# Completion of the DrugMatrix Toxicogenomics Database using ToxCompl

**DOI:** 10.1101/2024.03.26.586669

**Authors:** Guojing Cong, Robert M. Patton, Frank Chao, Daniel L. Svoboda, Warren M. Casey, Charles P. Schmitt, Charles Murphy, Jeremy N. Erickson, Parker Combs, Scott S. Auerbach

**Author notes:** Corresponding Authors: Guojing Cong Scott S. Auerbach.

## Abstract

The DrugMatrix Database contains systematically generated toxicogenomics data from short-term in vivo studies for over 600 chemicals. However, most of the potential endpoints in the database are missing due to a lack of experimental measurements. We present our study on leveraging matrix factorization and machine learning methods to predict the missing values in the DrugMatrix, which includes gene expression across eight tissues on two expression platforms along with paired clinical chemistry, hematology, and histopathology measurements. One major challenge we encounter is the skewed distribution of the available measured data, in terms of both tissue sources and values. We propose a method, ToxiCompl, that applies systematic hybrid sampling guided by Bayesian optimization in conjunction with low-rank matrix factorization to recover the missing values. ToxiCompl achieves good training and validation performance from a machine learning perspective.

We further conduct an in-depth validation of the predicted data from biological and toxicological perspectives with a series of analyses. These include examining the connectivity pattern of predicted gene expression responses, characterizing molecular pathway-level responses from sets of differentially expressed genes, evaluating known transcriptional biomarkers of tissue toxicity, and characterizing predicted apical endpoints. Our analysis shows that the predicted differential gene expression, broadly speaking, aligns with what would be anticipated. For example, in most instances, our predicted differentially expressed gene lists offer a connectivity level comparable to that of measured data in connectivity analysis. Using Havcr1, a known transcriptional biomarker of kidney injury, we identify treatments that, based on the predicted expression data, manifest kidney toxicity in a manner that is mechanistically plausible and supported by the literature. Characterization of the predicted clinical chemistry data suggests that strong effects are relatively reliably predicted, while more subtle effects pose a greater challenge. In the case of histopathological prediction, we find a significant overprediction due to positivity bias in the measured data. Developing methods to deal with this bias is one of the areas we plan to target for future improvement. The main advantage of the ToxiCompl approach is that, in the absence of additional experimental data, it drastically extends the toxicogenomic landscape into a number of data-poor tissues, thereby allowing researchers to formulate mechanistic hypotheses about effects in tissues that have been underrepresented in the literature. All measured and predicted DrugMatrix data (i.e., gene expression, clinical chemistry, hematology, and histopathology) are available to the public through an intuitive GUI interface that allows for data retrieval, gene set analysis and high dimensional visualization of gene expression similarity (https://rstudio.niehs.nih.gov/complete_drugmatrix/).

## 1 Introduction

Data is critical for modeling relationships between effects historically used to assess toxicity and those measured by newer approaches, such as toxicogenomics. A major challenge in developing new toxicity assessment methods is the limited availability of data. DrugMatrix represents a significant effort to create an integrated data resource, characterizing both traditional endpoints, e.g., histopathology/clinical pathology, and whole genome transcriptomics in the same animals. With over 600 chemicals studied in a collection of short-term in vivo rat studies, it is the largest systematically generated toxicogenomics database. As such, the data has been extensively reused in many studies (e.g. see [36]), particularly for deriving signatures of toxicity effects and mechanistic processes.

Two different rat microarray platforms were used to analyze gene expression from DrugMatrix samples: the GE Healthcare CodeLink UniSet Rat I (RU1) [2], a first-generation microarray technology no longer available, and the Affymetrix GeneChip Rat Genome 230 2.0 (RG230) [1], a second-generation technology still in use [101]. In DrugMatrix, the tissue/sample coverage is more comprehensive with the CodeLink platform, measuring data for eight tissues including liver, kidney, heart, skeletal muscle, spleen, bone marrow, intestine, and brain. In contrast, Affymetrix covers only liver, kidney, heart, and skeletal muscle. Most DrugMatrix microarray studies are paired with standard clinical pathology measurements and targeted histopathology (i.e., pathology performed on the same tissues as the transcriptomic measurements). Due to cost constraints, each study conducted to create the DrugMatrix resource was targeted based on prior knowledge of target organ toxicity. Consequently, many studies that make up the DrugMatrix data set are focused on the most common target organs, liver and kidney, with much more limited assessment of less common target tissues (e.g., spleen and brain). Therefore, when assembled in matrix form—studies (chemical-dose level-duration) by endpoints (tissue gene expression, tissue histopathology, clinical pathology)—the DrugMatrix data is missing about 88% of potential endpoints. These missing data are primarily gene expression endpoints in tissues not typically overtly effected by chemical exposure, as noted earlier. Here we apply artificial intelligence (AI) and machine learning methods to infer these missing endpoints.

In recent years, innovative AI methodologies have been employed to extend and create gene expression data, leveraging the power of covariance inference methods. For instance, techniques like L1000 and S1500+, in conjunction with the GENIE application, have shown promising results [100, 71, 72].These methods take subgenomic measurements and extrapolate them to whole-genome expression profiles, echoing the principles used in haplotype tagging single-nucleotide polymorphisms (SNPs) to infer complete genotypes in genomewide association studies (GWAS) [27]. This approach allows for a comprehensive understanding of genomic expressions by analyzing a fraction of the genome.

Recently, GAN-based methods developed at the National Center for Toxicological Research (NCTR), particularly ToxGAN and TransOrGAN [21, 66], have represented a significant advancement in inferring patterns of gene expression. These methods distinctively integrate various factors such as chemical structure, affected organ, dosage level, and exposure duration in ToxGAN, or age and inter-tissue correlative patterns of expression in TransOrGAN, to predict whole-genome expression. In addition, the NCTR group has also applied GAN technology in a similar manner to what was done with ToxGAN, creating AnimalGAN, which infers patterns of clinical pathological response to chemical challenges [20]. These AI-driven approaches demonstrate varying degrees of success and have the potential to closely approximate actual gene expression measurements. Collectively, these efforts highlight the inherent connectivity and covariant behavior of biological features across different levels of organization in complex biological systems. Moreover, they demonstrate the ability of AI-driven methods to identify and utilize these relationships to enhance the biological data landscape. In alignment with this conceptual framework, we describe a method that capitalizes on these relationships to infer the missing fraction of the DrugMatrix data.

The matrix form of the DrugMatrix dataset naturally lends itself to methods such as matrix imputation [117], completion [89, 55], and factorization [60]. There is a substantial body of literature on the theory and application of matrix methods in data science, with diverse applications including remote sensing [96], collaborative filtering [90], dimensionality reduction [97], biomedical link prediction [84], and machine learning classification [8].In our study, we adopt low-rank factorization to complete the DrugMatrix dataset. Initially, we establish that enough endpoints are present to apply the completion method. We then address the challenge posed by the extremely skewed data distribution in DrugMatrix for accurate prediction. A straightforward application of matrix completion results in the loss of rare but important signals. Practically all data in DrugMatrix, which are of interest to practitioners (e.g., over and under-expressed genes), are rare. To preserve these rare signals, we propose hybrid sampling methodologies based on the discrete form of the data derived from DrugMatrix. We have named our approach ToxiCompl. Our study demonstrates that ToxiCompl significantly enhances prediction performance for such signals in comparison to plain matrix factorization methods.

We validate the output of ToxiCompl in two distinct ways. The first method adheres to standard AI and machine learning practices, where we hold out a small portion of the data (e.g., 5%) from the original matrix and compute the deviation, for example, the mean absolute error (MAE) of the predicted data. We achieve an MAE of 0.09 and a mean *F*_1_ score (see definition in section 3) of 0.5 for our predictions. The second method involves investigating the predicted data from a biological and toxicological perspective. This includes studying the connectivity pattern of the predicted gene expression patterns, conducting pathway analysis to characterize molecular pathway-level responses from sets of differentially expressed genes, leveraging known transcriptional biomarkers of tissue toxicity for validation, and characterizing predicted apical endpoints. Our validation and analysis demonstrate the potential impact of the completed DrugMatrix on future toxicology studies. Given that our approach is purely computational and does not require further animal testing, its success heralds new research directions and could have a fundamental impact on future toxicogenomics studies.

The rest of the paper is organized as follows. In section 2, we present the details of DrugMatrix and introduce the factorization algorithm with initial results; in section 3 we focus on techniques to preserve rare signals while still maintaining a low prediction error; we validate the predicted matrix in section 5 through extensive analysis from a biological and toxicological perspective; in section 4 we discuss several potential alternative approaches for predicting the missing endpoints, and summarize some results of using these alternative approaches; we give our conclusion and future work in section 6.

## 2 Matrix Completion with DrugMatrix

Matrix completion recovers a matrix *M* of size *n*_1_ *×n*_2_ from a subset of its entries [22]. Obviously assumptions have to be made about the full matrix for successful recovery, and most studies assume it is low-rank. Low rank matrix completion is well-studied with many theoretic results and practical applications (including the famous Netflix challenge [29]). Matrix completion algorithms minimize the rank, the nuclear norm, or the Frobenius norm. For a survey, see [79]. Beyond plain completion, in real-world applications, oftentimes additional information is associated with the rows and columns. Take the Netflix challenge for example. The rows have associated user information, and the columns have associated movie information. Such information can be incorporated as additional features, and they in turn may be leveraged to improve completion performance (e.g., see [23, 69, 119, 81]) or to predict for new rows or columns where there are no existing entries at all. The latter is often referred as the cold-start problem (e.g., see [94, 12, 42, 118]) in the literature.

### 2.1 Statistics of DrugMatrix

The DrugMatrix data set when organized into a matrix format contains approximately *n*_1_ = 193, 000 rows and *n*_2_ = 3, 000 columns. The DrugMatrix data types in the matrix used in this study include histopathology (treatment group average severity score, 0-4; 146 metrics), clinical chemistry (treatment group average, value varies based on endpoint, 19 metrics), hematology (treatment group average, value varies based on endpoint, 19 metrics) and gene expression on Codelink and Affymetrix gene expression platforms (log10 ratio to a set of controls). Each column in the data set represents an individual treatment group (chemical-dose-duration) average for each endpoint. Out of a total of about 580M entries, close to 72M or 12.5% are present. Table 1 shows the distribution of data present for different expression platforms, RU1 and RG230, and organs, Liver (LI), Kidney (KI), Heart (HE), Bone Marrow (BM), Brain (BR), Intestine (IN), Spleen (SP), and Skeletal Muscle (SM). There are much more LI and KI data than the other types. There are much fewer BR, IN, and BM data. In fact, there are 100 times fewer IN data than LI data. The original data matrix along with all results of the study can be found here (https://cebs-ext.niehs.nih.gov/cebs/paper/15888/private/Mjc1MDIwNmU4NmQxMTNjMDhlYjdlMzA3NzUxMmI5NmUK).

**Table 1:** Data of different categories.

| **RU1** | **RG230** | **LI** | **KI** | **HE** | **BM** | **BR** | **IN** | **SP** | **SM** |
| --- | --- | --- | --- | --- | --- | --- | --- | --- | --- |
| 32.6M | 39.4M | 34.5M | 19.0M | 11.8M | 2.7M | 0.55M | 0.17M | 1.5M | 1.5M |

The RU1 and RG230 genomic data are,a s noted above, log10 ratio to control within the range of [-4.195, 3.878]. Within this range, depending on the expression level, the data fall in 5 categories: extremely under-expressed (-2), under expressed (1), normal (0), over expressed (1), and extremely over expressed (2). Table 2 shows the distribution of data into 5 different categories. Category 0 dominates the other categories, with about 92% of all available data samples.

**Table 2:** The distribution of the categories in the categorical data and predicted test data by the original model.

| **Category -2** | **Category -1** | **Category 0** | **Category 1** | **Category 2** |
| --- | --- | --- | --- | --- |
| 0.03% | 4.09% | 91.94% | 3.88% | 0.036% |

We first check the necessary conditions for applying matrix completion to generate plausible missing entries.

We establish that the ‘complete’ matrix would be low-rank by inspecting the organization of the matrix. The rank *r* of DrugMatrix shall obviously be no bigger than *n*_2_ *≈* 3, 000. In fact, we expect *r ≪ n*_2_ as the 3,000 treatments are for only 636 drugs, and for each drug, there are no more than 5 different treatment regimes. We also know some of these drugs should produce similar or correlated probes. Among the rows (*n*_1_ *≈* 193, 000), correlations exist between the corresponding RU1 and RG230 measurements for the same probes. In addition, the same probe exists for eight organs. Our analysis suggests that DrugMatrix is likely low-rank.

We then check whether there are enough data entries present to recover the whole matrix. Several results are developed with regard to the theoretical bounds of reconstruction. Keshavan, Montanari, and Oh [55] proposed an algorithm that reconstructs the matrix with *m* = *O*(*rn*_2_) entries with relative root mean square error

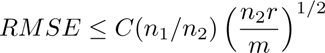

with *C* being a numerical constant. Cand’es and Tao [15] showed that *m ≥ Cµ*_0_*n*_1_*rlog*(*n*_1_) entries are necessary for completion when the entries are sampled uniformly at random. After estimating the constants *C* and *r*, we can verify that there are likely enough samples in DrugMatrix for low-rank approximation. Recht [89] improves on prior work, and with little assumptions on the matrix shows that *m ≥ max{µ*^2^*, µ*_0_*}r*(*n*_1_ + *n*_2_)*β* log^2^(2*n*_1_) is needed for reconstruction. Here *µ*_0_ and *µ*_1_ are some positives related to the bounds on the coherences and maximum entries of the matrix.

### 2.2 Completion

We adopt the popular Funk-SVD approach [86] for completion, where the *M* is factored into *P × Q*. Intuitively, *P* maps the probes into a *r* dimensional latent space, and *Q* maps the treatments into the same *r* dimensional latent space. *P* is of size *n*_1_ *× r*, and *Q* is of size *n*_2_ *× r*. This formulation is common in corroborative filtering applications such as the Netflix challenge [41]. *P* and *Q* are randomly initialized and then learned from the observed entries in the matrix, using the following objective function

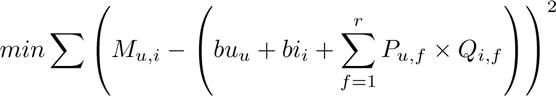

with *M_u,i_* being the observed entries, *bu_u_* and *bi_i_* being the biases for the probes and treatments, respectively. A solution to the problem can be found with stochastic optimization. We use two popular optimizers in our study: stochastic gradient descent (SGD) [91] and Adam [57].

In our initial experiment with training, we randomly partition the data into 90:5:5 train-validation-test split. Unless noted otherwise, when the optimizer is SGD, we use a learning rate of 0.01, weight decay of 0.01, and *r* = 300; when the optimizer is Adam, we use a learning rate of 0.001 with the same weight decay and *r*. The models are all trained for 300 epochs with early stopping.

Table 3 shows the performance measured by MAE, accuracy, and *F*_1_ score of matrix completion on DrugMatrix. Completion is performed for both regression and classification. In regression, the test MAE achieved is 0.09. The classification task predicts the categories ([-2, -1, 0, 1, 2]) for test entries. Completion achieves a test accuracy of 90.98%, with an *F*_1_ of 95.11%.

**Table 3:** Performance of completion.

| Test MAE | Accuracy (category) | $F_1$ |
| --- | --- | --- |
| 0.09 | 90.98% | 95.11% |

Figure 1 shows test MAE for the individual organs. For comparison purpose, we include the performance of a heuristic called Nearest-Neighbor-Mean (NNM), that for each entry to be predicted, picks *k* (e.g., in our study we use *k* = 10) nearest available entries in the same row and computes the mean.

**Figure 1:**
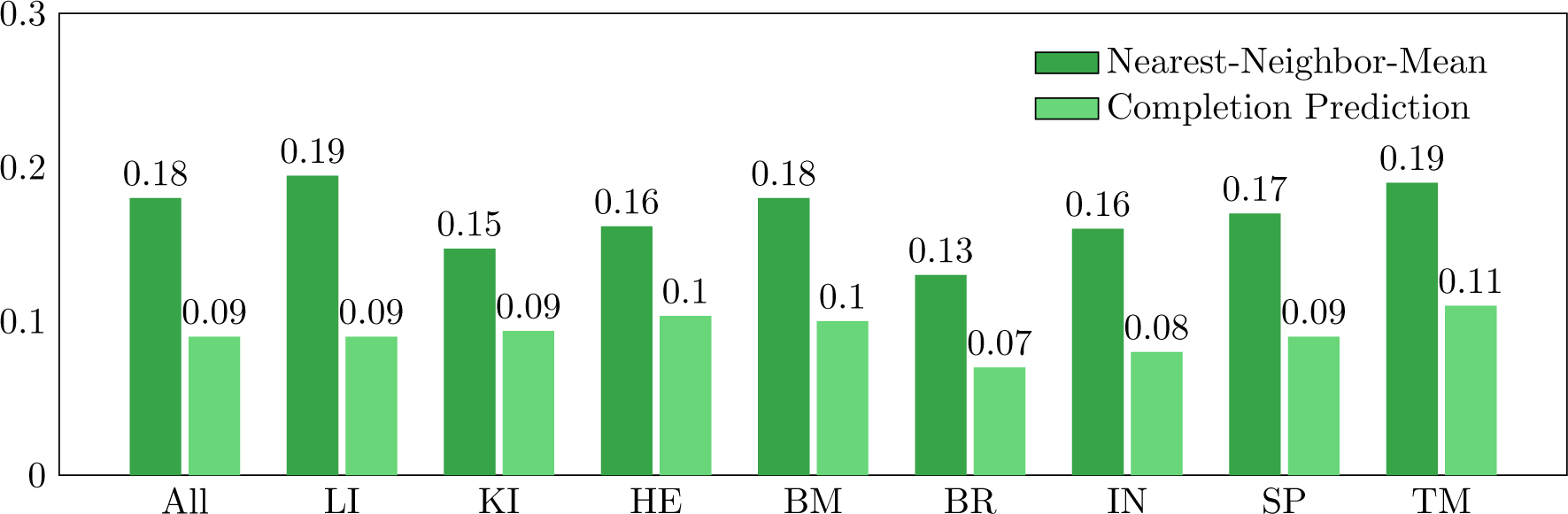
Matrix completion results for the overall DrugMatrix (All) and for each organs.

In Figure 1, it is clear that matrix completion beats NNM for regression for all organs, demonstrating its effectiveness.

## 3 Preserving Rare Signals

DrugMatrix carries meaning beyond just a low-rank matrix of numbers and its patterns have important biological implications. For example, positive values in the RU1 and RG230 data indicate over-expression while negative values indicate under-expression. The extreme values are of special interests to researchers and practitioners. The imbalance of distribution of data as shown in Table 2 and the potential usage of predicted data requires considerations beyond straightforward matrix completion.

Table 2 shows that most of the available entries in DrugMatrix fall within Category 0. Thus in practice, the other categories Categories -2, -1, 1, and 2 carry more useful information. From a toxicology perspective, we would want to recover these values as accurately as possible. Unfortunately, there are very few data samples available in these categories making it challenging to predict accurately for them. Only about 0.03% available data fall within Categories -2 and 2 each. They require special attention for prediction.

We investigate the performance of matrix completion for preserving rare signals. We compute *precision*, *recall*, and *F*_1_ *score* for category prediction, and the results on the test set are shown in Table 4. From Table 4 it is obvious that the prediction is not equally accurate for all categories. In fact, the performance for every category other than 0 is very poor, with *F*_1_ scores of 0.0%, 0.9%, 1.17%, and 0.0% for Categories -2, -1, 1, and 2, respectively.

**Table 4:** The precision, recall, and *F*_1_ score by category on test data.

| Categories | Precision | Recall | $F_1$ Score |
| --- | --- | --- | --- |
| -2 | 0.00% | 0.00% | 0.00% |
| -1 | 0.16% | 0.50% | 0.90% |
| 0 | 91.96% | 98.89% | 95.29% |
| 1 | 4.36% | 0.68% | 1.17% |
| 2 | 0.00% | 0.00% | 0.00% |

To account for the performance of preserving rare signals, we use the *mean of F*_1_ *score* (Mean *F*_1_) across all categories as our metric. Thus Mean *F*_1_=0.1948 for matrix completion. The reason for this behavior becomes apparent when we look at the distribution of data that matrix completion predicts. The results are shown in Table 5. In Table 5, almost 99% of the data points are predicted to be in Category 0. In comparison, only 0.0031% and 0.0025% are predicted for Category 2 and -2, respectively, while 0.4999% and 0.6075% are predicted for Category 1 and -1, respectively. Clearly due to the vast majority of training data in Category 0, matrix completion learns to predict most data entries to be in Category 0.

**Table 5:** The distribution of the categories in test data and predicted data.

| Categories | Ground Truth Distribution | Predicted Distribution |
| --- | --- | --- |
| -2 | 0.0328% | 0.0031% |
| -1 | 4.0970% | 0.4999% |
| 0 | 91.9464% | 98.8870% |
| 1 | 3.8880% | 0.6075% |
| 2 | 0.0359% | 0.0025% |

To improve the prediction performance for the rare categories, we augment matrix completion so that it gives more “weight” to data in categories other than 0. There are several ways to achieve this. We can augment the objective function to penalize heavily mis-predictions in the non-zero categories. We can also apply non-uniform sampling to the different categories. We find non-uniform sampling works better for completing DrugMatrix. For the rest of the paper, unless noted otherwise, our study with sampling operates on the DrugMatrix in its category form. That is, all entries are in [-2, -1, 0, 1, 2]. We will present results with DrugMatrix in its continuous form in Section 3.4

### 3.1 Over and Under Sampling

We first experiment to train the model with data that are uniformly drawn from all categories. Both random over-sampling and random under-sampling are considered. For over-sampling, data in the minority categories (all categories other than 0) are duplicated until the distribution is uniform. For under-sampling, data points are randomly removed from all of the categories except -2 until the distribution is uniform. Note that when under-sampling, a different sample is taken between each epoch during training, while during over-sampling a single over-sampled training data set is used for the entirety of the training period.

We again randomly partition the data into 90:5:5 train-validation-test split. During training, we either over or under sample, and run SGD for 300 epochs. The results in comparison to the baseline implementation (the straightforward completion) are shown in Figure 2.

**Figure 2:**
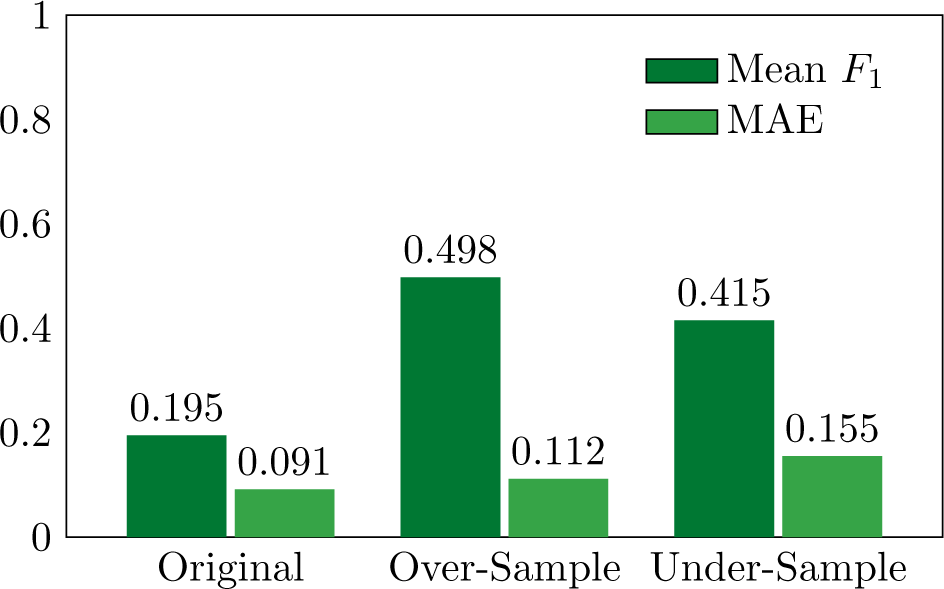
Mean *F*_1_ Scores and MAE for Different Sampling Distributions. Original denotes the baseline

As seen in Figure 2, both over-sampling and under-sampling substantially improve Mean *F*_1_ scores over the baseline. Both approaches perform much better at accurately predicting data for the minority categories. In fact, the improvement with Mean *F*_1_ over the baseline is about 2.5x and 2.1x for over-sampling and under-sampling, respectively. We note a slight degradation in MAE for both approaches. There appears to be a tug-of-war between Mean *F*_1_ and MAE. This unsurprisingly reflects the tension between high overall performance and high performance for under-represented categories in DrugMatrix.

### 3.2 Hybrid Sampling

We propose a hybrid sampling approach to further improve upon the results from over-sampling and undersampling. The goal is to improve Mean *F*_1_ while minimizing MAE. In hybrid sampling, we under-sample the majority category while simultaneously over-sample the minority categories. Hybrid sampling can be considered as a superset of strategies over under-sampling and over-sampling. We experiment different sampling distributions, as shown in Table 6.

**Table 6:**
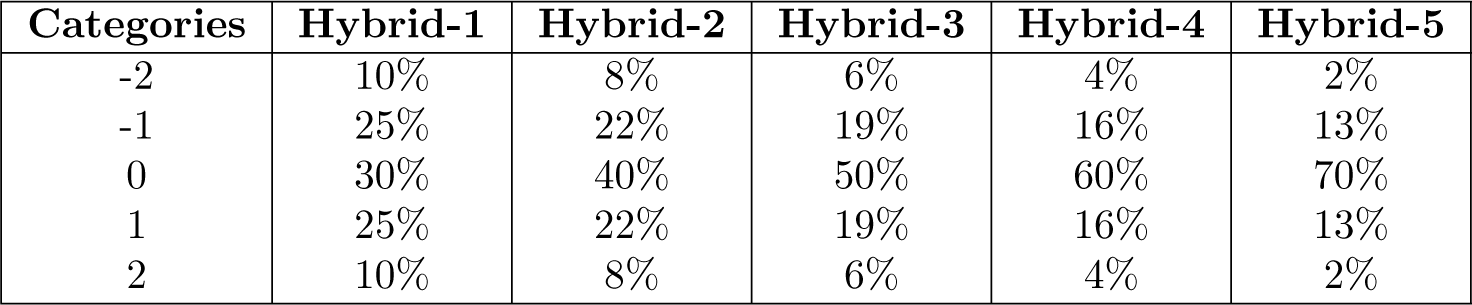
Distributions tried for hybrid sampling.

| Categories | Hybrid-1 | Hybrid-2 | Hybrid-3 | Hybrid-4 | Hybrid-5 |
| --- | --- | --- | --- | --- | --- |
| -2 | 10% | 8% | 6% | 4% | 2% |
| -1 | 25% | 22% | 19% | 16% | 13% |
| 0 | 30% | 40% | 50% | 60% | 70% |
| 1 | 25% | 22% | 19% | 16% | 13% |
| 2 | 10% | 8% | 6% | 4% | 2% |

The five different distributions in Table 6 are evaluated in our experiments. In training we use the same hyper-parameters as before and train for 300 epochs. Of all distributions, Hybrid-5 yields the lowest MAE and highest Mean *F*_1_ score, as seen in Figure 3. In fact, of all the implementations, Hybrid-5 has the highest Mean *F*_1_ with a MAE slightly bigger than baseline. The Mean *F*_1_ is 2.61x better than baseline.

**Figure 3:**
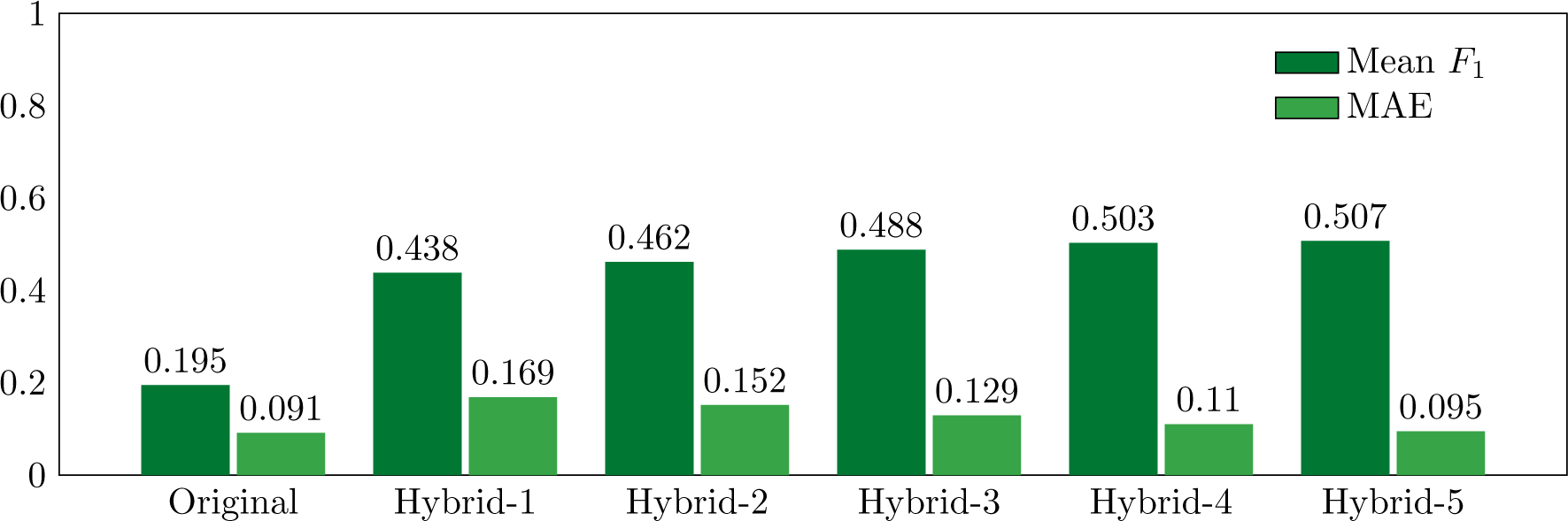
Mean *F*_1_ scores and MAE on test data with different hybrid sampling implementations.

### 3.3 Bayesian Optimization for finding the optimal distribution

The distributions of samples chosen from different categories so far are chosen manually for hybrid sampling. We apply Bayesian optimization to find “optimal distributions” for the best performance. Bayesian optimization is a global optimization technique for black box functions. Here the inputs are the distributions and the optimization target is the Mean *F*_1_ score. We design our Bayesian optimization approach to find the optimal sampling distribution to maximize the Mean *F*_1_ score.

We start with 10 initial distributions. Since Hybrid-5 yields good performance already, we include Hybrid-5 within the 10 and randomly select 9 other distributions. Our objective is to maximize the Mean *F*_1_ score by finding the right distribution. We use Scikit-Learn and experiment with three acquisitions functions (expected improvement, probability of improvement, and lowest confidence bound). We also explore representing the distribution in three different ways. In the first, for each category the input is between 0 and 1 and all of the inputs are normalized to calculate the distribution. In the second, the input is the ratio for each of the minority categories and the remainder is used as ratio for the majority category. In the third, for each category the inputs are the number of data points to include.

The best distribution produced by Bayesian optimization is shown in Table 7 with a Mean *F*_1_ of 0.5122 and MAE of 0.0916. The Mean *F*_1_ is about 2.63x better than baseline and the MAE is comparable to baseline.

**Table 7:** The distribution generated by Bayesian optimization with the maximal Mean *F*_1_.

| Categories | Distribution |
| --- | --- |
| -2 | 4.3229% |
| -1 | 10.6839% |
| 0 | 67.0960% |
| 1 | 12.8972% |
| 2 | 5.0000% |

### 3.4 Leveraging Category Data to Predict Continuous DrugMatrix

We first summarize the performance of our implementations on category data in Figure 4. Both MAE and Mean *F*_1_ score are shown. In Figure 4, the various sampling approaches combined with matrix completion achieve substantially better Mean *F*_1_ scores than baseline matrix completion and NNM. The baseline implementation achieves the best MAE. Only Hybrid-5 and Bayesian optimization are able to also achieve a MAE comparable to baseline.

**Figure 4:**
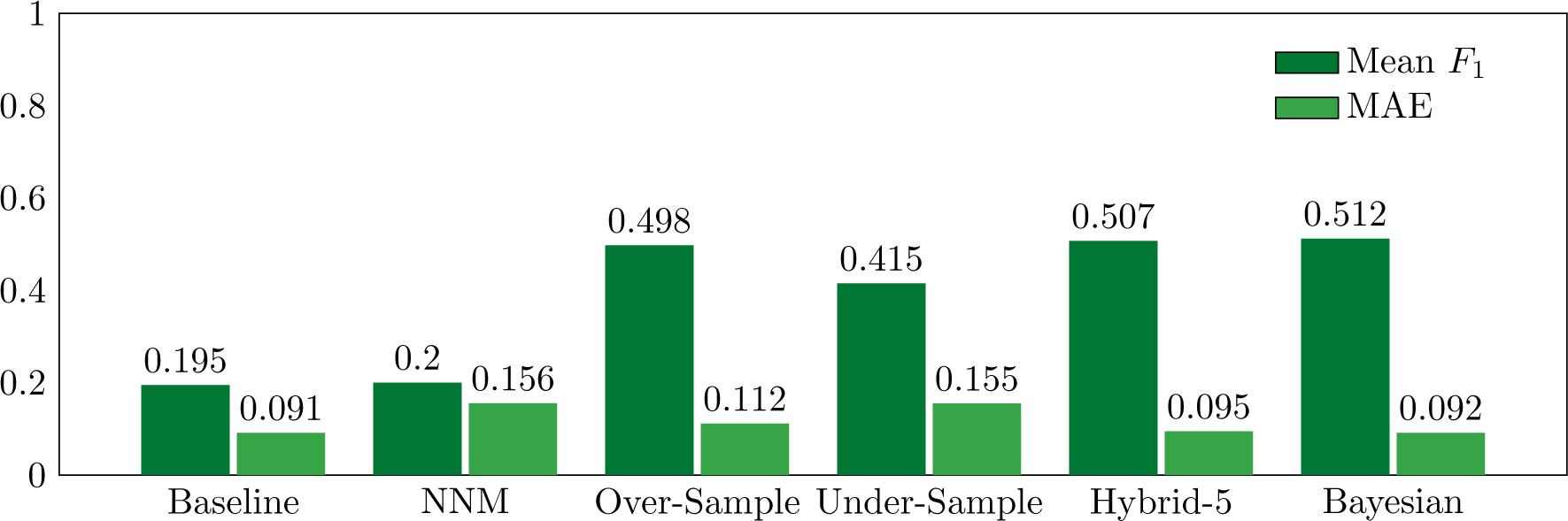
Mean *F*_1_ Scores and MAE for Hybrid Sampling Distributions.

We leverage the category data for regression prediction on the continuous DrugMatrix in order to retain information present in the minority categories. We create a combined data set that include both the category and continuous values for each data sample. The category values are used to sample the distribution when matrix completion is applied to train on the continuous data sample. We use *Bayesian optimization* as our hybrid sampling strategy. We call our low-rank factorization for DrugMatrix with sampling frequencies optimized by Bayesian optimization ToxiCompl, and its detailed mechanism is shown in Figure 5.

**Figure 5:**
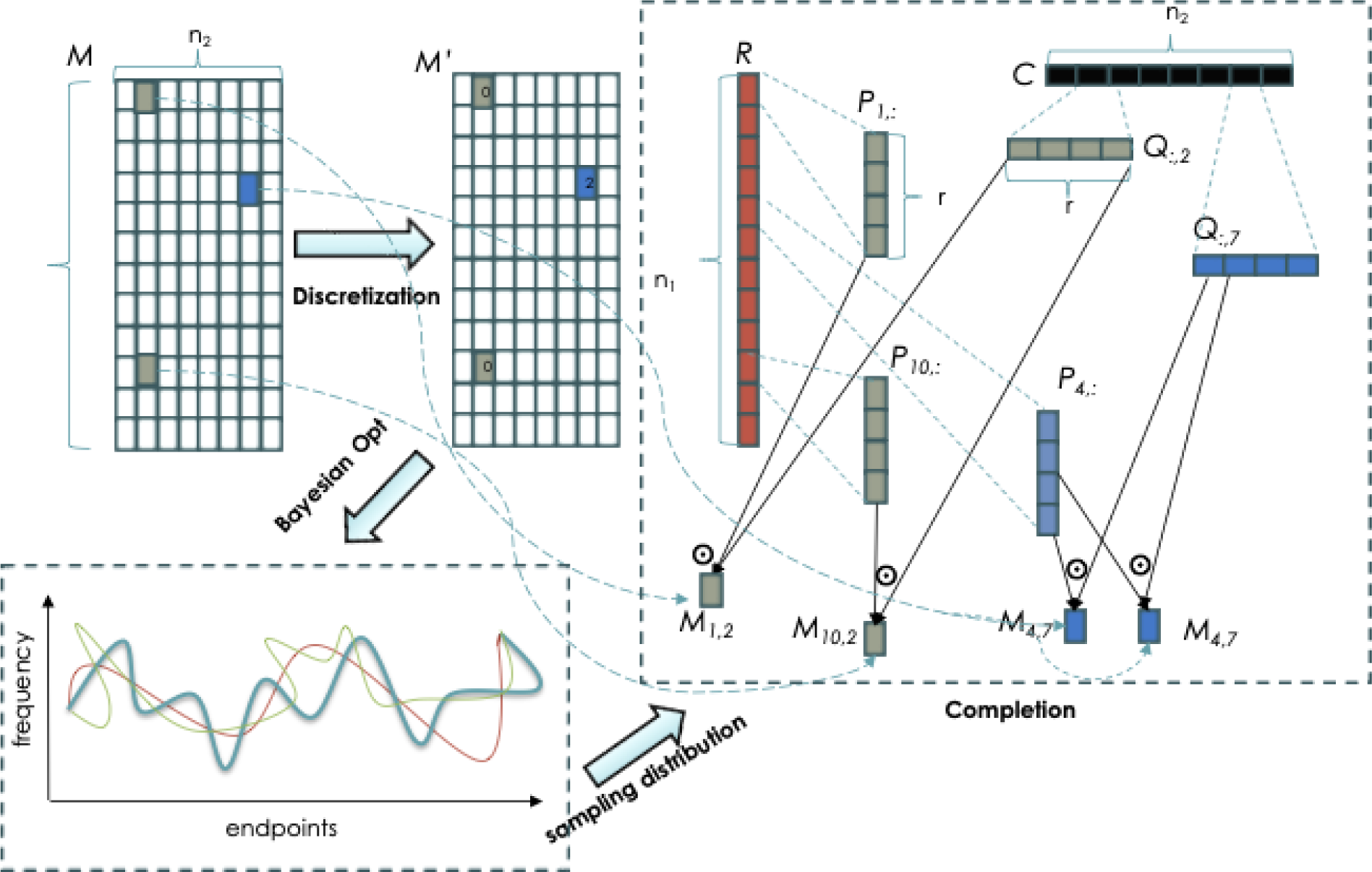
ToxiCompl components with control and data flow.

In Figure 5, the original matrix is shown as *M*. As an example, we show a 12*x*6 matrix, with *M*_1,2_, *M*_4,7_, and *M*_10,2_ containing measured values. All other entries are missing. ToxiCompl attempts to recover all these values while preserving the rare signals (colored blue). ToxiCompl first disretize *M* into *M ^′^* with category values. In our example, two values are in the 0 category while the remaining one is in category 2. Matrix *M ^′^* is an input to the Bayesian optimizer so that it can compute a sampling distribution of *M* for the “completion” component to learn the two low-rank matrices, *P_i,_*: and *Q*:,_*j*_. The Bayesian optimizer observes the validation performance of the “completion” component relative to the sampling distribution of *M*, and produces an optimal distribution. In the figure, *M*_4,7_ is sampled twice (in one training epoch) due to the relative rarity of 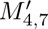 in *M*^′^, while *M*_1,2_ and *M*_10,2_ are only sampled once. The “completion” component computes the sampled *M* entries by 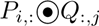, here 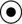 denotes dot product (for simplicity and readability, we do not include the biases for each rows and columns in the example).

We compare the performance of ToxiCompl to both the baseline and NNM on the original DrugMatrix data. The results on overall MAE and MAE by category are shown in Figure 6. The overall MAE of *Bayesian optimization* is 0.094, similar to that of Baseline, 0.091, and about two times smaller than that of NNM, 0.180. For the underrepresented categories, *Bayesian optimization* performs better than Baseline, and on average about two times better than NNM.

**Figure 6:**
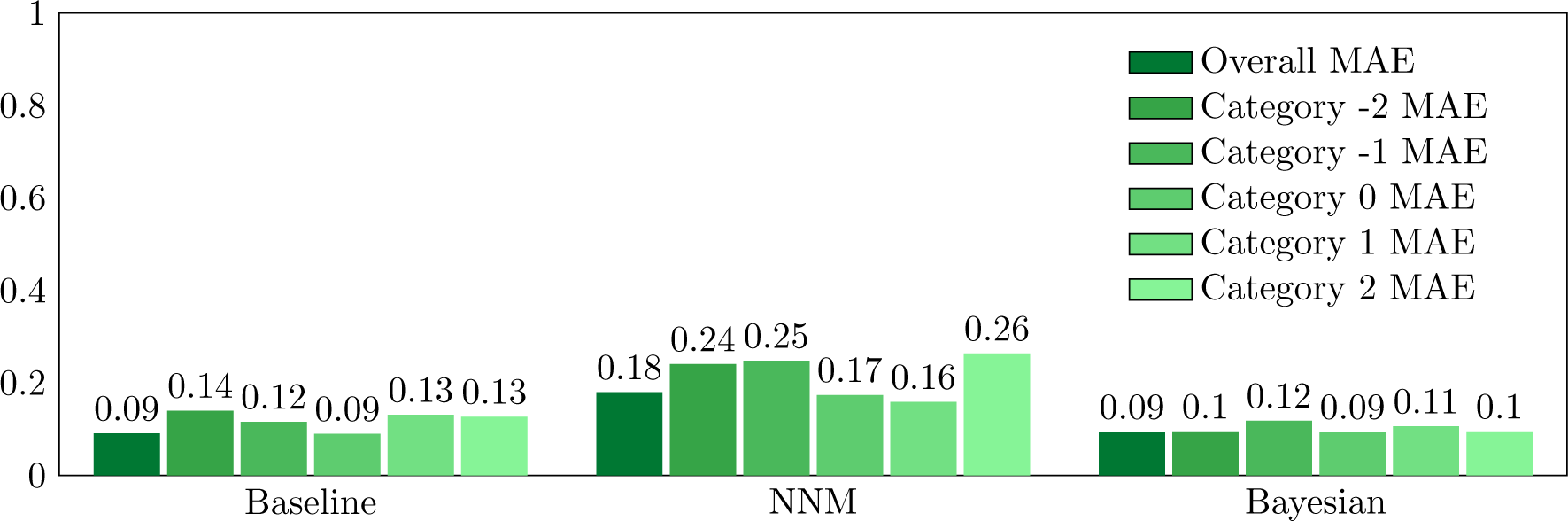
Overall MAE and MAE by category for continuous test data.

## 4 Alternative Approaches

DrugMatrix contains a rich collection of auxiliary data such as dosage, treatment duration, and gene marker that enable other approaches than ToxiCompl to fill the missing entries. Rather than modeling the interaction between treatments and responses as a matrix, we can also construct a graph to model the relationship and apply graph learning. We discuss and explore some of these possibilities for comparison purpose.

Predicting missing entries in DrugMatrix can be cast as an inference problem after a machine learning model is trained on the entries present. However, this is quite different from the regular machine learning setup where for example a 8:1:1 train-validation-test split, is typically used. We need to define inputs or features for machine learning models, and these can not be the entries in the Matrix. We extract three features for each row in DrugMatrix. They are *Organ*, *Platform*, and *Marker*. *Organ* takes one of eight distinct values: LI, KI, HE, BM, BR, IN, SP, and SM. *Platform* is either RU1 or RG230. There are however 34021 different *Marker* values. Three features can be extracted from the columns. They are *Drug*, *Dosage*, and *Duration*. *Drug* has 636 values, *Dosage* has 339 values, and *Duration* has 10 values.

### 4.1 Random Forest

We train a random forest regressor with the entries in the DrugMatrix. Each of the input contain 6 features, *Organ*, *Platform*, *Marker*, and *Drug*, *Dosage*, and *Duration*. Our experiments achieve train MAEs between 0.08 and 0.11, and test MAEs around 0.16, when we vary the number of tree estimators and the max depth of the trees. The performance of random forest is inferior to ToxiCompl.

Random forest does reveal the importance of the features for learning. With 100 tree estimators, the feature importance is shown in Figure 7.

**Figure 7:**
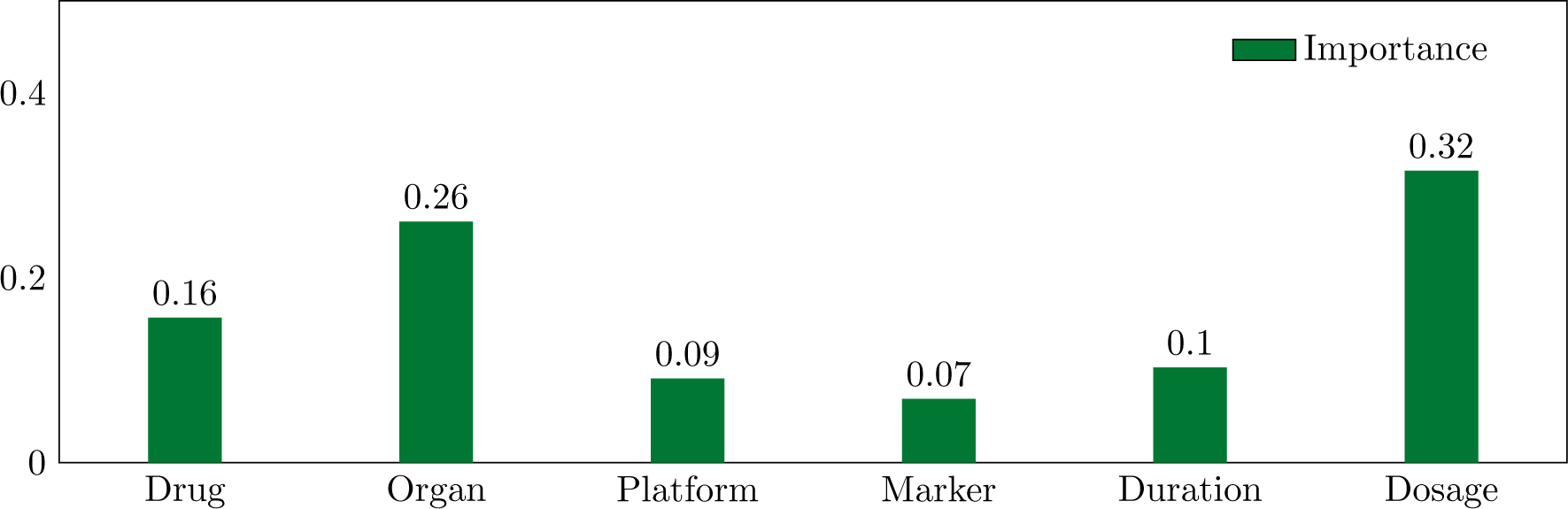
Feature importance for trained random forest regressor.

Figure 7 shows that *Drug*, *Organ*, and *Dosage* are much more important than *Platform*, *Marker*, and *Duration*, although none of them are negligible. For this specific regressor, *Dosage* appears to be the most distinguishing feature, followed by *Organ*, and *Drug*.

### 4.2 Deep Neural Networks

We train a deep neural network regressor for predicting the missing entries in DrugMatrix. The network contains several fully connected layers each followed with an activation function (e.g., ReLU). We use the same six features as in the random forest regressor described in Section 4.1, and experiment with different number of layers (i.e., 2, 4, and 8 hidden layers) and different number of neurons per layer (e.g., 128, 256, and 512). The performance of all these experiments are surprisingly consistent, achieving test accuracies in the range [0.115, 0.116]. We note that deep neural networks perform worse than ToxiCompl, but better than random forest.

### 4.3 Completion with Side Features

Matrix completion itself can be augmented with side features. Intuitively, knowing the side feature values, for example, a specific dosage, may well be helpful to predict a gene marker response. Side features are associated with both the rows and columns, and for DrugMatrix, we have at least three side features, *Platform*, *Marker*, and *Duration*, for each row and three side features, *Drug*, *Organ*, and *Dosage*, for each column. We embed each feature (row and column) into a *r* dimension vector (*R* and *C*, respectively). Again, we construct *P* and *Q* to optimize the following objective function

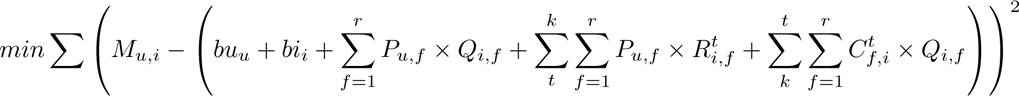

with *M_u,i_* being the observed entries, *bu_u_* and *bi_i_* being the biases for the gene expressions and the treatments, respectively. Here we have *k* row features, and *k* column features.

We compare matrix completion with and without side features, and the results is shown in Figure 8.

**Figure 8:**
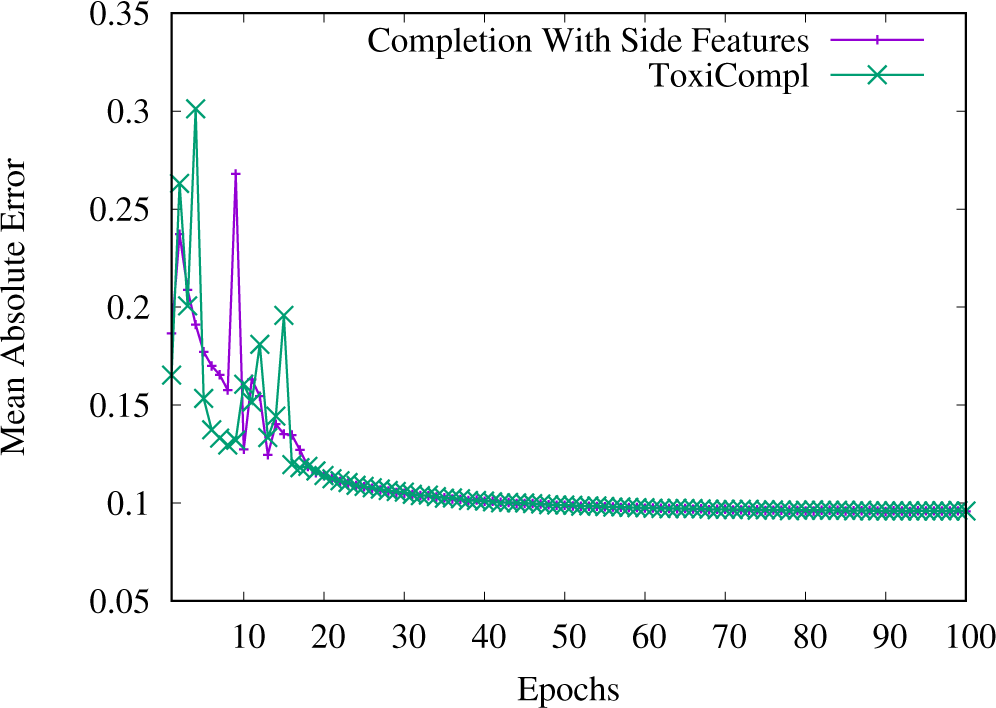
ToxiCompl vs. completion with side features.

In Figure 8, we plot the test MAE after each epoch. It appears that completion with side features shows more stable behavior than without side features. However, as training progresses, the two methods show similar performance. Deep neural network and matrix factorization can be further combined and be used in a scenario for predicting the responses for a completely new treatment without any measured data at all. We will explore this aspect in our future work.

### 4.4 Graph Neural Networks (GNNs)

We can construct a graph to model the interaction between the toxicogenomics profiles and treatments by creating nodes representing these entities and edges representing the responses, and apply graph learning to predict the missing entries in DrugMatrix.

There is a large body of work on applying GNNs to collaborative filtering (for a comprehensive survey, see [37]). Compared with matrix factorization, one of the advantages of GNNs is that it can explore higherorder connectivity than the first-order connectivity (straightforward similarity between users and items). In addition to the extremely skewed data distribution, applying GNNs in the toxicogenomics context presents unique computing challenges due to the extremely high degrees of the nodes (for treatments with measurement data present, the node degree is in the range of 32,000). We leave further exploration of GNNs to our future work.

## 5 Validation and Analysis

To assess the extent to which predicted data mirrors what would be anticipated from measured data, several biological interpretation techniques were utilized. In this assessment, emphasis was placed on evaluating predicted data without corresponding measured data. This approach ensures the elimination of potential information leakage from the measured data through cross-validation.

The initial method we employed for biological characterization is connectivity. In this method, predicted data from prototype agents undergoes correlative analysis with a broad range of signatures from the Drug-Matrix database. The goal is to determine if the predicted expression aligns globally with treatments from similar prototype agents.

In our second evaluation, we investigated if there is biologically coherent pathway activation across tissues evident in the predicted gene expression from the prototype agent, Fenofibrate.

For the third assessment, we executed paired comparisons of pathway enrichment between closely matched predicted and measured data across various tissues. The objective was to see if the top enriched pathways were consistent between both sets of data.

In the fourth evaluation, we analyzed the plausibility and validity of toxicity-related transcriptional biomarker induction observed in the predicted data across several tissues.

Lastly, we conducted a review of the predicted apical data, performing a plausibility evaluation of predicted findings in comparison to closely related treatments that had measured clinical chemistry and histopathology results.

### 5.1 Connectivity Analysis

Connectivity is a metric for determining the degree of similarity between lists of differentially expressed genes [95]. The stronger the connectivity the greater the similarity in the up and down regulated genes between two lists of differentially expressed genes. Connectivity is often used to match patterns produced by prototype agents with those of unknown mechanism of action to infer the mechanistic effects of the unknown. To evaluate the accuracy of the predicted gene expression, 5 gene lists derived from treatments with prototype agents (i.e., they have known mechanisms of action associate with specific patterns of transcriptomic response) were selected from the RG230 data that was solely predicted (i.e., there was not any corresponding measured gene expression data). This analysis is limited to liver gene expression because there are well defined mechanisms of action in this tissue [109]. The top 500 genes as determined by absolute value of fold change for each of these treatments was loaded into the Illumina Correlation Engine and connectivity analysis was performed with the 3-day Affymetrix DrugMatrix Liver data. The statistical metric used to quantify connectivity in the Illumina Correlation Engine is the Running Fischer test [62].

The first predicted treatment evaluated for connectivity was beta-naphthoflavone at a dose of 80 mg/kg for 28 days. Beta-naphthoflavone is an Ahr agonist that strongly up-regulates Cyp1a1, Cyp1a2, and Cyp1b1 [82]. This prediction of this treatment indicates that these genes would be up-regulated following this treatment (see Complete DrugMatrix application). Table 8 below presents the top treatments with the greatest connectivity. Results in the table represent connectivity as quantified by the Running Fischer test p-value, direction of correlation, and the number of overlapping genes between the predicted treatment and the measured DrugMatrix data.

**Table 8:** Connectivity analysis to measured data – beta-naphthoflavone.

| Bioset Name | P-value | Correlation / Direction | Common Genes |
| --- | --- | --- | --- |
| Liver of rats + NIMESULIDE at 162mg-kg-d in corn oil by oral gavage 3d _vs_ vehicle | 7.40E-39 | + | 121 |
| Liver of rats + LEFLUNOMIDE at 60mg-kg-d in corn oil by oral gavage 3d _vs_ vehicle | 1.10E-34 | + | 94 |
| Liver of rats + CARVEDILOL at 2000mg-kg-d in corn oil by oral gavage 3d _vs_ vehicle | 3.00E-33 | + | 121 |
| Liver of rats + LOMEFLOXACIN at 2000mg-kg-d in CMC by oral gavage 3d _vs_ vehicle | 2.00E-32 | + | 101 |
| Liver of rats + AMIODARONE at 147mg-kg-d in CMC by oral gavage 3d _vs_ vehicle | 1.00E-31 | + | 95 |

Interestingly, among the top 5 most connected treatments, only leflunomide is an AhR activator. A review of the connecting genes showed that the connectivity of the four non-AhR activating treatments was primarily through down-regulated genes. This suggests two possibilities: 1) There might be a secondary mechanism, separate from AhR activation, influencing gene expression in both beta-naphthoflavone and the four connective non-AhR treatments. 2) The predicted expression for beta-naphthoflavone might be inaccurate, at least for the down-regulated genes. This implies that there could be only partial accuracy in the predicted expression for beta-naphthoflavone.

The next predicted gene list that was evaluated for connectivity was from primidone. Primidone is metabolized in vivo to phenobarbital [14], which activates the nuclear receptors CAR and PXR. This activation leads to an increased expression of Cyp2b1, Cyp2b2, and Cyp3a9 in the liver [74]. The predicted gene expression response from a treatment of Primidone at a dose of 750 mg/kg for 5 days, indicated an increased expression of all three of these genes (see Complete DrugMatrix application). Connectivity analysis using the top 500 predicted genes by fold change revealed the strongest connectivity with the treatments listed in

**Table 9:** Connectivity analysis to measured data – primidone.

| Bioset Name | P-value | Correlation / Direction | Common Genes |
| --- | --- | --- | --- |
| Liver of rats + ANASTROZOLE at 400mg-kg-d in CMC by oral gavage 3d _vs_ vehicle | 7.50E-34 | + | 175 |
| Liver of rats + CLONAZEPAM at 2500mg-kg-d in CMC by oral gavage 3d _vs_ vehicle | 1.50E-25 | + | 115 |
| Liver of rats + CARBAMAZEPINE at 490mg-kg-d in CMC by oral gavage 3d _vs_ vehicle | 2.10E-24 | + | 109 |
| Liver of rats + METHAPYRILENE at 100mg-kg-d in CMC by oral gavage 3d _vs_ vehicle | 2.80E-23 | + | 149 |
| Liver of rats + AMINOGLUTETHIMIDE at 350mg-kg-d in CMC by oral gavage 3d _vs_ vehicle | 3.30E-23 | + | 116 |

All these compounds are documented activators of CAR/PXR [54] or show strong induction of Cyp2b1, Cyp2b2, and Cyp3a9 consistent with activation of these receptors (observed in Illumina Correlation Engine). In the case primidone the predicted data show appropriate connectivity.

Clofibrate is a well-known activator of the nuclear receptor PPAR*α*, which is used therapeutically to treat hyperlipidemia [30]. PPAR*α* activators lead to an increased expression of many genes in the rat liver associated with fatty acid metabolism [103] Some of the most pronounced transcriptionally responsive genes to PPAR*α* activation include Acot1, Cyp4a1, and Vnn1. The treatment with Clofibrate at a dose of 500 mg/kg for 3 days was not measured using Affymetrix microarrays as part of the DrugMatrix Database. However, data for this treatment were predicted as part of this project. As anticiapted, Acot1, Cyp4a1, and Vnn1 were predicted to be up-regulated in the Clofibrate-500 mg/kg-3-day treatment (see Complete DrugMatrix application). The treatments most closely connected to the Clofibrate-500 mg/kg-3-day regimen are listed in Table 10.

**Table 10:** Connectivity analysis to measured data – Clofibrate.

| Bioset Name | P-value | Correlation / Direction | Common Genes |
| --- | --- | --- | --- |
| Liver of rats + NAFENOPIN at 338mg-kg-d in corn oil by oral gavage 3d _vs_ vehicle | 1.60E-77 | + | 198 |
| Liver of rats + BEZAFIBRATE at 617mg-kg-d in corn oil by oral gavage 3d _vs_ vehicle | 6.60E-75 | + | 172 |
| Liver of rats + FENOFIBRATE at 215mg-kg-d in corn oil by oral gavage 3d _vs_ vehicle | 7.10E-71 | + | 182 |
| Liver of rats + PIRINIXIC ACID at 364mg-kg-d in CMC by oral gavage 3d _vs_ vehicle | 2.10E-67 | + | 187 |
| Liver of rats + CLOFIBRIC ACID at 448mg-kg-d in corn oil by oral gavage 3d _vs_ vehicle | 1.40E-64 | + | 154 |

The top five connected treatments are all well-known PPAR*α* activators [103]. These findings clearly demonstrate that the predicted gene expression data yield expected patterns of gene expression, exhibiting robust connectivity to mechanistically relevant measured treatments.

Beta-estradiol is a pharmacological estrogen known for producing a distinct gene expression pattern in the male rat liver. Genes commonly up-regulated in the liver by such estrogenic treatments include Rbp7, Cited4, and Lifr (observed in the Illumina Correlation Engine). The predicted expression response from the treatment with beta-estradiol 3-benzoate at a dose of 25 mg/kg for 5 days showed up-regulation of Rbp7, Cited4, and Lifr suggesting the model can accurately predict response to this estrogenic treatment (see Complete DrugMatrix application).

Connectivity analysis btween the predicted gene expression response of beta-estradiol 3-benzoate-25mg/kg-5day treatment and the measured DrugMatrix samples yielded expected results, showing connectivity with measured estrogenic treatments (Table 11). An exception was the connectivity observed with dexamethasone, which is structurally related to estrogens but is a glucocorticoid receptor agonist. The results of the connectivity analysis largely indicate that the predicted gene expression is consistent with the effects of an estrogen in the liver.

**Table 11:**
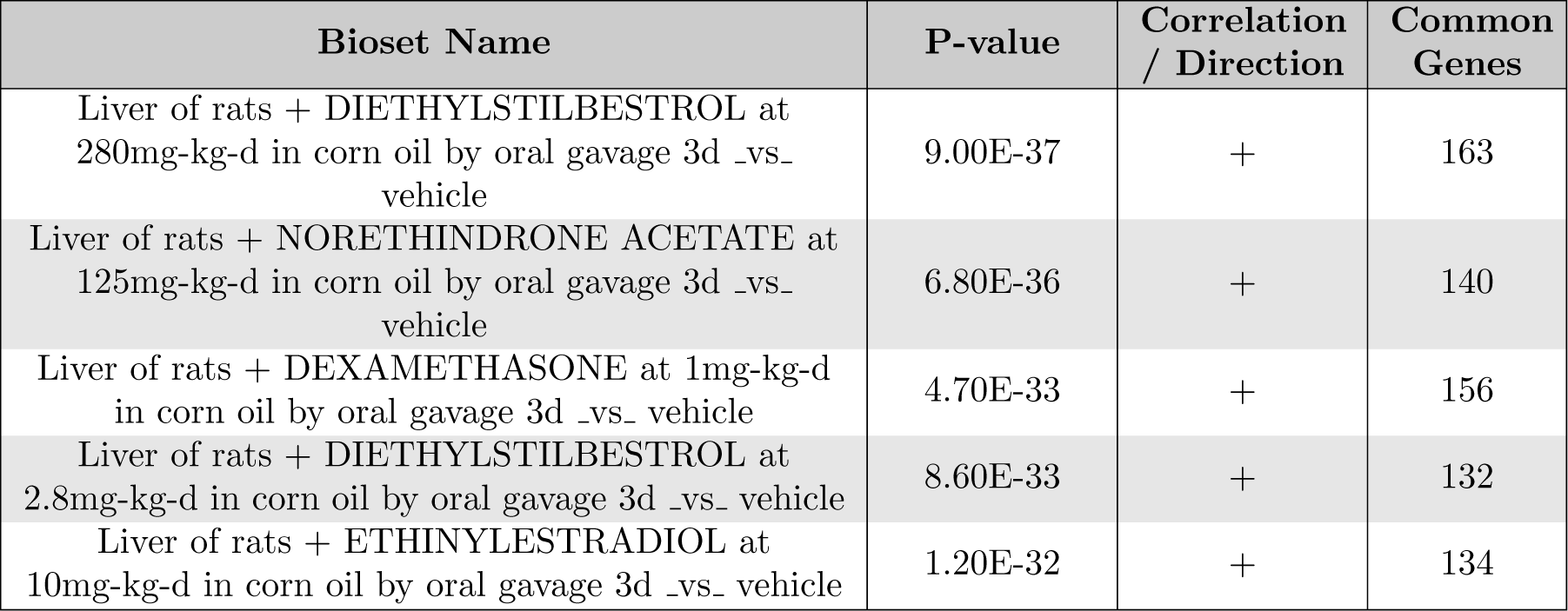
Connectivity analysis to measured data – beta-estradiol.

| Bioset Name | P-value | Correlation / Direction | Common Genes |
| --- | --- | --- | --- |
| Liver of rats + DIETHYLSTILBESTROL at 280mg-kg-d in corn oil by oral gavage 3d _vs_ vehicle | 9.00E-37 | + | 163 |
| Liver of rats + NORETHINDRONE ACETATE at 125mg-kg-d in corn oil by oral gavage 3d _vs_ vehicle | 6.80E-36 | + | 140 |
| Liver of rats + DEXAMETHASONE at 1mg-kg-d in corn oil by oral gavage 3d _vs_ vehicle | 4.70E-33 | + | 156 |
| Liver of rats + DIETHYLSTILBESTROL at 2.8mg-kg-d in corn oil by oral gavage 3d _vs_ vehicle | 8.60E-33 | + | 132 |
| Liver of rats + ETHINYLESTRADIOL at 10mg-kg-d in corn oil by oral gavage 3d _vs_ vehicle | 1.20E-32 | + | 134 |

N-nitrosodiethylamine (NDEA) is a member of the highly genotoxic and carcinogenic class of compounds commonly referred to as nitrosamines [65]. NDEA induces an increase in a specific set of liver genes indicative of a genotoxic effect [32]. Among these genes are Ccng1, Phlda3, Aen, Mdm2, and Cdkn1a, all integral to the P53 signaling network [3]. Liver gene expression data using Affymetrix microarrays for the treatment N-nitrosodiethylamine at a dose of 34 mg/kg over 5 days was not produced as part of the original DrugMatrix dataset. However, the predicted Affymetrix data for this treatment showed an up-regulation of all five aforementioned genes (see Complete DrugMatrix application), therefore aligning with the expected response in the P53 pathway following a high dose exposure to N-nitrosodiethylamine. The connectivity analysis, presented in Table 12, using the predicted data revealed the highest scoring connectivity with treatments known for their well-documented genotoxic properties [59, 43].

**Table 12:** Connectivity analysis to measured data – N-nitrosodiethylamine.

| Bioset Name | P-value | Correlation / Direction | Common Genes |
| --- | --- | --- | --- |
| Liver of rats + N-NITROSODIETHYLAMINE at 34mg-kg-d in saline by oral gavage 3d _vs_ vehicle | 2.60E-31 | + | 100 |
| Liver of rats + STAVUDINE at 1400mg-kg-d in water by oral gavage 3d _vs_ vehicle | 5.40E-29 | + | 142 |
| Liver of rats + AFLATOXIN B1 at .3mg-kg-d in CMC by oral gavage 3d _vs_ vehicle | 6.80E-23 | + | 84 |
| Liver of rats + 2-ACETYLAMINOFLUORENE at 30mg-kg-d in CMC by oral gavage 3d _vs_ vehicle | 3.60E-21 | + | 100 |
| Liver of rats + MITOMYCIN C at 1.7mg-kg-d in saline by intravenous 3d _vs_ vehicle | 1.90E-19 | + | 67 |

Overall, the findings from the connectivity analysis suggest that the predicted expression, broadly speaking, aligns with what would be anticipated. In most instances, this prediction offers a connectivity level comparable to that of observed data. However, there’s an essential caveat to this assertion: our focus was on the liver due to its well-defined modes of action. This type of detailed analysis isn’t currently feasible for other organs. As a result, the validity of connectivity based on predicted gene expression in other tissues might not achieve the same accuracy level observed in the liver.

### 5.2 Pathway Analysis – Fenofibrate Across All Tissues

Pathway analysis is a means of characterizing molecular pathway-level responses from sets of differentially expressed genes [80]. Enriched pathways provide higher-order insights into the biological processes altered by a treatment. An example of this is the impact of fibrate drugs on genes within various fatty acid metabolism pathways [83]. In rats, due to the somewhat ubiquitous expression of PPAR*α* across most tissues evaluated in DrugMatrix, it is anticipated that most tissues would respond to fibrate exposure, primarily through the up-regulation of fatty acid metabolism [33]. To evaluate the predictive data for its capacity to identify pathways of known effect, we used the RU1 predicted data (i.e., treatments for which there were no existing measured data) from fenofibrate treatments and checked for up-regulation in pathways related to fatty acid metabolism. For this analysis, we limited our enrichment to genes that were up-regulated by more than 1.5-fold. The results are shown in Table 13

**Table 13:**
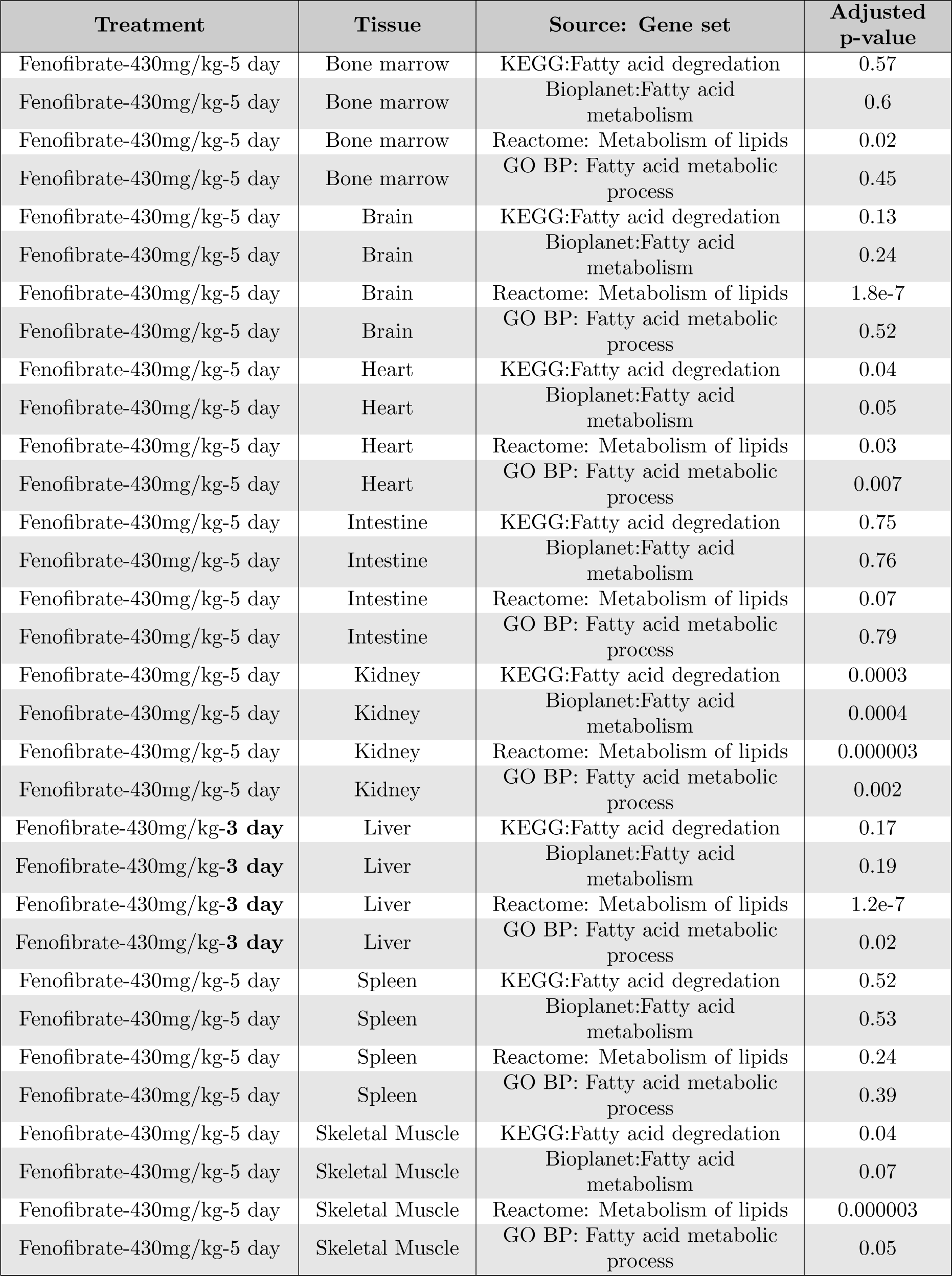
PPAR*α* associated pathway enrichment across tiss – Fenofibrate.

| Treatment | Tissue | Source: Gene set | Adjusted p-value |
| --- | --- | --- | --- |
| Fenofibrate-430mg/kg-5 day | Bone marrow | KEGG:Fatty acid degradation | 0.57 |
| Fenofibrate-430mg/kg-5 day | Bone marrow | Bioplanet:Fatty acid metabolism | 0.6 |
| Fenofibrate-430mg/kg-5 day | Bone marrow | Reactome: Metabolism of lipids | 0.02 |
| Fenofibrate-430mg/kg-5 day | Bone marrow | GO BP: Fatty acid metabolic process | 0.45 |
| Fenofibrate-430mg/kg-5 day | Brain | KEGG:Fatty acid degradation | 0.13 |
| Fenofibrate-430mg/kg-5 day | Brain | Bioplanet:Fatty acid metabolism | 0.24 |
| Fenofibrate-430mg/kg-5 day | Brain | Reactome: Metabolism of lipids | 1.8e-7 |
| Fenofibrate-430mg/kg-5 day | Brain | GO BP: Fatty acid metabolic process | 0.52 |
| Fenofibrate-430mg/kg-5 day | Heart | KEGG:Fatty acid degradation | 0.04 |
| Fenofibrate-430mg/kg-5 day | Heart | Bioplanet:Fatty acid metabolism | 0.05 |
| Fenofibrate-430mg/kg-5 day | Heart | Reactome: Metabolism of lipids | 0.03 |
| Fenofibrate-430mg/kg-5 day | Heart | GO BP: Fatty acid metabolic process | 0.007 |
| Fenofibrate-430mg/kg-5 day | Intestine | KEGG:Fatty acid degradation | 0.75 |
| Fenofibrate-430mg/kg-5 day | Intestine | Bioplanet:Fatty acid metabolism | 0.76 |
| Fenofibrate-430mg/kg-5 day | Intestine | Reactome: Metabolism of lipids | 0.07 |
| Fenofibrate-430mg/kg-5 day | Intestine | GO BP: Fatty acid metabolic process | 0.79 |
| Fenofibrate-430mg/kg-5 day | Kidney | KEGG:Fatty acid degradation | 0.0003 |
| Fenofibrate-430mg/kg-5 day | Kidney | Bioplanet:Fatty acid metabolism | 0.0004 |
| Fenofibrate-430mg/kg-5 day | Kidney | Reactome: Metabolism of lipids | 0.000003 |
| Fenofibrate-430mg/kg-5 day | Kidney | GO BP: Fatty acid metabolic process | 0.002 |
| Fenofibrate-430mg/kg-**3 day** | Liver | KEGG:Fatty acid degradation | 0.17 |
| Fenofibrate-430mg/kg-**3 day** | Liver | Bioplanet:Fatty acid metabolism | 0.19 |
| Fenofibrate-430mg/kg-**3 day** | Liver | Reactome: Metabolism of lipids | 1.2e-7 |
| Fenofibrate-430mg/kg-**3 day** | Liver | GO BP: Fatty acid metabolic process | 0.02 |
| Fenofibrate-430mg/kg-5 day | Spleen | KEGG:Fatty acid degradation | 0.52 |
| Fenofibrate-430mg/kg-5 day | Spleen | Bioplanet:Fatty acid metabolism | 0.53 |
| Fenofibrate-430mg/kg-5 day | Spleen | Reactome: Metabolism of lipids | 0.24 |
| Fenofibrate-430mg/kg-5 day | Spleen | GO BP: Fatty acid metabolic process | 0.39 |
| Fenofibrate-430mg/kg-5 day | Skeletal Muscle | KEGG:Fatty acid degradation | 0.04 |
| Fenofibrate-430mg/kg-5 day | Skeletal Muscle | Bioplanet:Fatty acid metabolism | 0.07 |
| Fenofibrate-430mg/kg-5 day | Skeletal Muscle | Reactome: Metabolism of lipids | 0.000003 |
| Fenofibrate-430mg/kg-5 day | Skeletal Muscle | GO BP: Fatty acid metabolic process | 0.05 |

Overall, the results from the pathway analysis were underwhelming. We anticipated consistent upregulation of the chosen pathways/gene sets across all tissues, but this was not observed. Perhaps the most surprising outcome was the marginal findings in the liver, where these gene sets are known to respond robustly to PPAR*α* activation.

### 5.3 Pathway Analysis – Top 10 Pathways Measured vs. Predicted across Different Tissues

Ifosfamide is a genotoxic chemotherapy drug used for cancer treatment. It adversely impacts bone marrow by suppressing blood cell production, which can lead to anemia, decreased white blood cell counts, and reduced platelet counts. To assess the validity of the pathways identified from the predicted bone marrow gene data (RU1), we compared the treatments: ifosfamide-143mg/kg-5day (measured) and ifosfamide-143mg/kg-3day (predicted, with no corresponding measured data). These treatments are expected to have largely similar pathway activation profiles. For this analysis, we concentrated on the up-regulated genes and utilized the Enrichr tool for pathway enrichment evaluation. The top 10 enriched KEGG pathways in bone marrow gene expression from both treatments are presented in Table 14.

**Table 14:** Top enriched KEGG pathways measured vs predicted – Ifosfamide.

|  | Ifosfamide-143mg/kg-5d<br>(measured) |  | Ifosfamide-143mg/kg-3d<br>(predicted) |  |
| --- | --- | --- | --- | --- |
| Replicated<br>in predicted<br>data | KEGG pathway | Adjusted<br>p-value | KEGG pathway | Adjusted<br>p-value |
| X | Pathways in cancer | 2.93E-12 | Pathways in cancer | 4.84E-09 |
| X | Rap1 signaling<br>pathway | 8.09E-09 | Rap1 signaling<br>pathway | 1.65E-07 |
|  | PI3K-Akt signaling<br>pathway | 1.35E-07 | TGF-beta signaling<br>pathway | 1.52E-06 |
| X | Proteoglycans in<br>cancer | 1.35E-07 | Hippo signaling<br>pathway | 2.79E-05 |
| X | Calcium signaling<br>pathway | 2.55E-07 | Proteoglycans in<br>cancer | 2.79E-05 |
| X | TGF-beta signaling<br>pathway | 8.86E-07 | MAPK signaling<br>pathway | 2.95E-05 |
|  | Apelin signaling<br>pathway | 8.86E-07 | Focal adhesion | 3.43E-05 |
|  | AGE-RAGE signaling<br>pathway in diabetic<br>complications | 9.22E-07 | Fluid shear stress and<br>atherosclerosis | 4.31E-05 |
| X | Focal adhesion | 5.09E-06 | Calcium signaling<br>pathway | 4.31E-05 |
| | Human papillomavirus<br>infection | 5.09E-06 | PPARα signaling<br>pathway | 4.31E-05 |

A striking overlap was observed in the top 10 enriched pathways from both the measured and predicted data, including the top 2 pathways on both lists. This finding suggests that, at the pathway level, the predicted bone marrow expression data aligns with what would likely be observed if the predicted data were empirically characterized.

In the heart, Angiotensin II induces hypertrophy, fibrosis, and adverse remodeling [64]. To assess the validity of the pathways identified from the predicted heart gene expression data (RG230), we compared the treatments: angiotensin II-4.2mg/kg-5day (measured) and angiotensin II-15mg/kg-5day (predicted, with no corresponding measured data). These treatments are expected to exhibit largely similar pathway activation profiles. For this analysis, we focused on the up-regulated genes and utilized the Enrichr tool to evaluate pathway enrichment. The top 10 enriched KEGG pathways in heart gene expression from the treatments are presented in Table 15.

**Table 15:**
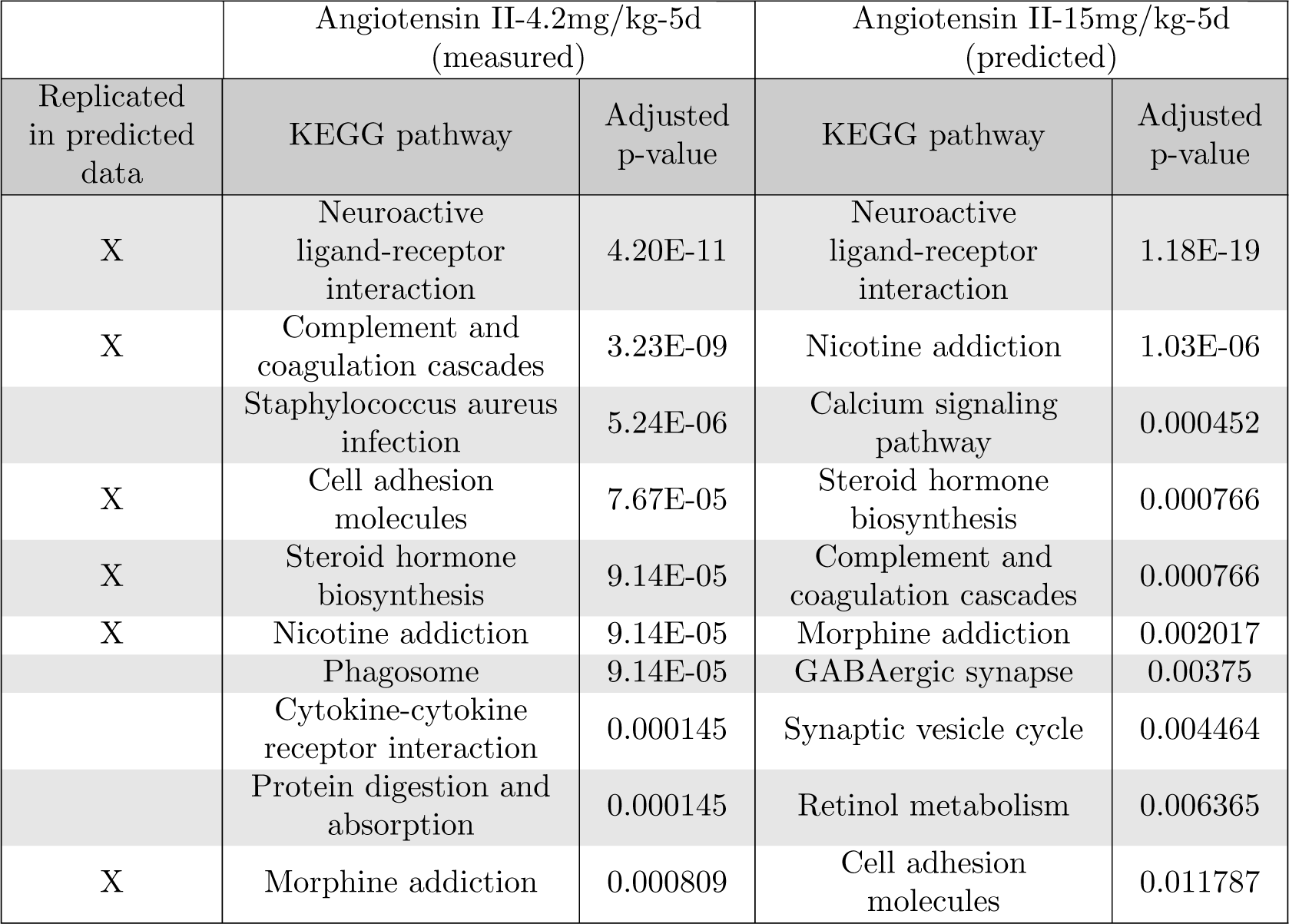
Top enriched KEGG pathways measured vs predicted – Angiotensin.

|  | Angiotensin II-4.2mg/kg-5d<br>(measured) |  | Angiotensin II-15mg/kg-5d<br>(predicted) |  |
| --- | --- | --- | --- | --- |
| Replicated<br>in predicted<br>data | KEGG pathway | Adjusted<br>p-value | KEGG pathway | Adjusted<br>p-value |
| X | Neuroactive<br>ligand-receptor<br>interaction | 4.20E-11 | Neuroactive<br>ligand-receptor<br>interaction | 1.18E-19 |
| X | Complement and<br>coagulation cascades | 3.23E-09 | Nicotine addiction | 1.03E-06 |
|  | Staphylococcus aureus<br>infection | 5.24E-06 | Calcium signaling<br>pathway | 0.000452 |
| X | Cell adhesion<br>molecules | 7.67E-05 | Steroid hormone<br>biosynthesis | 0.000766 |
| X | Steroid hormone<br>biosynthesis | 9.14E-05 | Complement and<br>coagulation cascades | 0.000766 |
| X | Nicotine addiction | 9.14E-05 | Morphine addiction | 0.002017 |
|  | Phagosome | 9.14E-05 | GABAergic synapse | 0.00375 |
|  | Cytokine-cytokine<br>receptor interaction | 0.000145 | Synaptic vesicle cycle | 0.004464 |
|  | Protein digestion and<br>absorption | 0.000145 | Retinol metabolism | 0.006365 |
| X | Morphine addiction | 0.000809 | Cell adhesion<br>molecules | 0.011787 |

Similar to the bone marrow gene expression analysis (see Table 14) a striking overlap was observed in the top 10 enriched pathways from both the measured and predicted data, including the top pathway on both lists. This finding suggests that, at the pathway level, the predicted heart gene expression data aligns with what would likely be observed if the predicted data were empirically characterized.

Clofibrate is a hypolipidemic drug that elicits various effects on the livers of rats. These effects include changes in enzyme expression, hepatocyte morphology, lipid composition, and increased peroxisome activity [63]. To assess the validity of the pathways identified from the predicted liver gene expression data (RG230), we compared the treatments: clofibrate-500mg/kg-5 day (measured) and clofibrate-500mg/kg-3 day (predicted, with no corresponding measured data). These treatments are expected to exhibit largely similar pathway activation profiles. For this analysis, we concentrated on the up-regulated genes and utilized the Enrichr tool for pathway enrichment evaluation. The top 10 enriched KEGG pathways in liver gene expression from these treatments are presented in Table 16.

**Table 16:** Top enriched KEGG pathways measured vs predicted – clofibrate-500 mg/kg/day.

|  | Clofibrate-500mg/kg-5d<br>(measured) |  | Clofibrate-500mg/kg-3d<br>(predicted) |  |
| --- | --- | --- | --- | --- |
| Replicated<br>in predicted<br>data | KEGG pathway | Adjusted<br>p-value | KEGG pathway | Adjusted<br>p-value |
| X | Neuroactive<br>ligand-receptor<br>interaction | 1.37E-12 | Fatty acid degradation | 4.46E-17 |
| X | Metabolism of<br>xenobiotics by<br>cytochrome P450 | 1.58E-11 | Neuroactive<br>ligand-receptor<br>interaction | 2.47E-14 |
| X | Peroxisome | 3.54E-11 | Peroxisome | 4.10E-14 |
| X | Fatty acid degradation | 4.75E-10 | PPAR signaling<br>pathway | 4.36E-14 |
| X | PPAR signaling<br>pathway | 5.00E-10 | Butanoate metabolism | 1.34E-05 |
| X | Steroid hormone<br>biosynthesis | 8.13E-10 | Retinol metabolism | 1.34E-05 |
|  | Valine, leucine and<br>isoleucine degradation | 7.03E-09 | Nicotine addiction | 3.62E-05 |
|  | Propanoate<br>metabolism | 2.10E-07 | Fatty acid elongation | 4.72E-05 |
|  | Drug metabolism | 2.11E-07 | Metabolism of<br>xenobiotics by<br>cytochrome P450 | 6.86E-05 |
| X | Retinol metabolism | 3.59E-07 | Steroid hormone<br>biosynthesis | 0.000118 |

As with the other pathway analysis results mentioned above, the liver gene expression reveals a significant overlap in the top 10 enriched pathways from both the measured and predicted data, including the top pathway on both lists. This finding suggests that, at the pathway level, the predicted liver expression data aligns with what would likely be observed if the predicted data were empirically characterized. Cisplatin is a chemotherapeutic agent that can induce damage to spleen tissue, trigger pro-inflammatory responses, and lead to structural changes in the spleen, including size reduction and hemosiderin accumulation [110]. To assess the validity of the pathways identified from the predicted spleen gene expression data (RU1), we compared the treatments: cisplatin-1.17mg/kg-3 day (measured) and clofibrate-1.17mg/kg-5 day (predicted, with no corresponding measured data). These treatments are expected to exhibit largely similar pathway activation profiles. For this analysis, we concentrated on the up-regulated genes and utilized the Enrichr tool for pathway enrichment evaluation. The top 10 enriched KEGG pathways in spleen gene expression from these treatments are presented in Table 17.

**Table 17:** Top enriched KEGG pathways measured vs predicted – Cisplatin-1.17 mg/kg/day.

|  | Cisplatin-1.17-3d (measured) |  | Cisplatin-1.17-5d (predicted) |  |
| --- | --- | --- | --- | --- |
| Replicated<br>in predicted<br>data | KEGG pathway | Adjusted<br>p-value | KEGG pathway | Adjusted<br>p-value |
|  | Calcium signaling pathway | 1.78E-05 | cAMP signaling pathway | 3.05E-05 |
| X | Complement and coagulation cascades | 0.000114143 | Pathways in cancer | 0.000358 |
| X | Pathways in cancer | 0.000133943 | Pertussis | 0.000802 |
|  | PI3K-Akt signaling pathway | 0.000235787 | AGE-RAGE signaling pathway in diabetic complications | 0.00121 |
|  | Choline metabolism in cancer | 0.000235787 | Fluid shear stress and atherosclerosis | 0.00121 |
|  | Endometrial cancer | 0.000850314 | beta-Alanine metabolism | 0.002408 |
|  | ErbB signaling pathway | 0.001998159 | TNF signaling pathway | 0.002408 |
|  | JAK-STAT signaling pathway | 0.003916979 | Cytokine-cytokine receptor interaction | 0.002519 |
| X | beta-Alanine metabolism | 0.008460076 | Rap1 signaling pathway | 0.004088 |

The pathway-level findings in the spleen were less consistent than in other tissues where this comparison was made. Only three out of the top 10 pathways from the measured data appeared in the top 10 pathways from the predicted data. These results indicate that the pathway insights derived from the predicted spleen gene expression data might be less reliable compared to other tissues.

Enalapril is an angiotensin-converting enzyme (ACE) inhibitor that is used to control blood pressure and proteinuria in the hypertensive patients. High doses can reduce renal blood flow, which lead to kidney damage [78]. To evaluate the validity of the pathways identified from predicted kidney gene expression data (RG230) we compared the following treatments enalapril-1500mg/kg-5 day (measured) to enalapril-1500mg/kg-3 day(predicted with no corresponding measured data). These treatments should have largely similar pathway activation profiles. For this analysis we focused on the up-regulated genes and used the Enrichr tool to evaluate pathway enrichment. The top 10 enriched KEGG pathways in spleen gene expression from treatments are shown in Table 18.

**Table 18:** Top enriched KEGG pathways measured vs predicted –Enalapril-1500 mg/kg/day.

|  | Enalapril-1500mg/kg-5d<br>(measured) |  | Enalapril-1500mg/kg-3d<br>(predicted) |  |
| --- | --- | --- | --- | --- |
| Replicated<br>in predicted<br>data | KEGG pathway | Adjusted<br>p-value | KEGG pathway | Adjusted<br>p-value |
| X | Neuroactive<br>ligand-receptor<br>interaction | 5.66E-09 | Neuroactive<br>ligand-receptor<br>interaction | 4.96E-27 |
| X | Metabolism of<br>xenobiotics by<br>cytochrome P450 | 4.96E-08 | Bile secretion | 6.21E-10 |
|  | Pentose and<br>glucuronate<br>interconversions | 1.03E-07 | Calcium signaling<br>pathway | 1.04E-09 |
| X | Chemical<br>carcinogenesis | 1.03E-07 | Steroid hormone<br>biosynthesis | 7.50E-09 |
|  | Glutathione<br>metabolism | 3.27E-07 | Metabolism of<br>xenobiotics by<br>cytochrome P450 | 9.17E-09 |
| X | Calcium signaling<br>pathway | 1.99E-06 | Nicotine addiction | 3.51E-07 |
| X | Steroid hormone<br>biosynthesis | 2.60E-05 | Retinol metabolism | 3.60E-07 |
|  | PPAR signaling<br>pathway | 2.60E-05 | Drug metabolism | 2.29E-06 |
| X | Drug metabolism | 2.60E-05 | Morphine addiction | 2.29E-06 |
|  | Ascorbate and<br>aldarate metabolism | 3.63E-05 | Chemical<br>carcinogenesis | 3.44E-06 |

Similar to many of the other pathway-level findings, there was a notable overlap in the top 10 enriched pathways from kidney data between both the measured and predicted datasets, including the top pathway on both lists. This finding suggests that, at a pathway level, the predicted kidney expression data aligns with what would be observed if the predicted data were empirically characterized.

Lovastatin is and HMG-CoA reductase inhibitor used treat hypercholesterolemia [44]. At high doses in rat it causes rhabdomyolysis (muscle cell lysis) which can lead to kidney damage. To evaluate the validity of the pathways identified from predicted skeletal muscle gene expression data (RU1) we compared the following treatments lovastatin-1500mg/kg-5 day (measured) to lovastatin-1500mg/kg-3 day (predicted with no corresponding measured data). These treatments should have largely similar pathway activation profiles. For this analysis we focused on the up-regulated genes and used the Enrichr tool to evaluate pathway enrichment. The top 10 enriched KEGG pathways in spleen gene expression from treatments are shown Table 19.

**Table 19:** Top enriched KEGG pathways measured vs predicted – Lovastatin-1500 mg/kg/day.

|  | Lovastatin-1500mg/kg-3d<br>(measured) |  | Lovastatin-1500mg/kg-5d<br>(predicted) |  |
| --- | --- | --- | --- | --- |
| Replicated<br>in predicted<br>data | KEGG pathway | Adjusted<br>p-value | KEGG pathway | Adjusted<br>p-value |
| X | Proteasome | 4.01E-13 | Proteasome | 0.007215 |
|  | Spinocerebellar ataxia | 2.33E-08 | Nucleotide excision<br>repair | 0.007215 |
| X | Valine, leucine and<br>isoleucine degradation | 8.57E-05 | Calcium signaling<br>pathway | 0.007215 |
| X | Drug metabolism | 9.03E-05 | PPAR signaling<br>pathway | 0.007215 |
|  | Fat digestion and<br>absorption | 0.000848 | Spinocerebellar ataxia | 0.017333 |
|  | Parkinson disease | 0.000859 | Valine, leucine and<br>isoleucine degradation | 0.021242 |
| X | PPAR signaling<br>pathway | 0.000859 | Cell cycle | 0.043375 |
|  | Epstein-Barr virus<br>infection | 0.001272 | Ubiquitin mediated<br>proteolysis | 0.049389 |
|  | Peroxisome | 0.002052 | Drug metabolism | 0.049389 |
| X | HIF-1 signaling<br>pathway | 0.002791 | HIF-1 signaling<br>pathway | 0.049389 |

Five out of the top 10 enriched pathways the measured and predicted lovastatin treatments in from skeletal muscle overlapped with the top pathway, proteosome, being the same for both data sets. This finding suggests the at a pathway level the predicted skeletal expression data is for most part consistent with what would be observed if the predicted data was empirically characterized.

The brain and intestine, which were not subjected to a measured vs. predicted pathway analysis in this section, were omitted because high-confidence pairings (i.e., high certainty of similar effects) could not be identified.

### 5.4 Transcriptional Biomarkers of Tissue Toxicity

To determine if the predicted data produced the expected transcriptional biomarker activation across various target tissues, we examined the purely predicted data for up-regulation of these genes in several tissues. The validity of the findings was assessed based on existing literature or by reviewing histopathological changes observed in DrugMatrix. All findings from this analysis including references to histopathological change can be found in the Shiny application described in section 5.6 that hosts the data from this study.

#### 5.4.1 Bone Marrow

Cdkn1a is a known transcriptional biomarker for bone marrow toxicity that acts through a genotoxic mechanism [17, 107]). In DrugMatrix, only the RU1 platform was used to measure gene expression in bone marrow; hence, the only predicted gene expression is from the RU1. A review of the measured Cdkn1a bone marrow gene expression data reveals that treatments that most strongly up-regulate Cdkn1a are toxic to the bone marrow and are typically genotoxic chemicals. Such treatments include doxifluridine, doxorubicin, ifosfamide, cyclophosphamide, and daunorubicin. In the predicted data for the corresponding measured treatments, all showed up-regulation of Cdkn1a (see Complete DrugMatrix application). This suggests that the inference approach performs as expected when training data are excluded from the model and then predicted.

For the data that was predicted without corresponding measured data, the following treatments were anticipated to elicit the greatest up-regulation of Cdkn1a: Cyclopropane carboxylic acid, doxorubicin (with a different duration and dose than the measured data), etoposide, gatifloxicin, and indarubicin. All of these compounds, with the exception of gatifloxicin, are well-documented genotoxic chemicals that cause bone marrow suppression. Overall, these findings suggest a moderate to high reliability for using solely predicted data to identify treatments that cause bone marrow suppression through a genotoxic mechanism.

#### 5.4.2 Brain

Spp1 is a recognized transcriptional biomarker that is up-regulated following overt neurotoxicity and traumatic brain injury in rats [99]. In DrugMatrix, only the RU1 platform was used to measure brain gene expression, meaning the only predicted gene expression derives from RU1. Reviewing the measured data, several well-known neurotoxic chemicals were found to up-regulate Spp1. These include high doses of caffeine [108], nicotine [34], haloperidol [34], and thimerosal [24]. In the predicted data corresponding to these treatments, all showed up-regulation of Spp1, indicating that the inference approach functions as anticipated when training data are omitted from the model and subsequently predicted (see Complete DrugMatrix application).

For the data predicted without corresponding measured data, the treatments predicted to cause the greatest up-regulation of Spp1 were progesterone, chlorambucil, meloxicam, hydralazine, and sodium nitroprusside (see Complete DrugMatrix application). Among these, only chlorambucil [48] and sodium nitroprusside [47] have documented neurotoxic effects. In conclusion, these results suggest limited reliability in using solely predicted data to identify neurotoxic treatments.

#### 5.4.3 Heart

Myh7 is a transcriptional biomarker that is up-regulated in the heart following a toxic challenge, which elicits histopathological changes such as single cell necrosis [76]. In DrugMatrix, Myh7 is only measured using the Affymetrix RG230 array, so the metrics for this analysis are derived from either measured or predicted Affymetrix data. A review of the measured heart Myh7 expression data identified treatments that exhibited the strongest up-regulation as sulindac, doxorubicin, dexamethasone, phenytoin, and cyclosporin A (see Complete DrugMatrix application). Doxorubicin, dexamethasone, and cyclosporin A are recognized cardiotoxic agents [77]. Sulindac, in the context of DrugMatrix, likely caused GI perforation, leading to high levels of systemic inflammation which may have induced a secondary stress on the heart, resulting in up-regulation of Myh7 [50]. Although phenytoin is not widely recognized as a cardiotoxic agent, it can induce cardiotoxicity at high doses through sodium channel inhibition [16].

Predicted expression levels for these treatments were consistent when Myh7 expression was excluded from the model and subsequently predicted. When evaluating the expression of Myh7 in treatments without corresponding measured data, the chemicals showing the highest up-regulation of the gene were miconazole, erythromycin, 1-napthyl isocyanate, benoxaprofen, and chlorpromazine (see Complete DrugMatrix application). Of these, only chlorpromazine is a well-documented cardiotoxicant [67]. In the DrugMatrix dataset, 1-napthyl isocyanate did result in an increase in relative heart weight, suggesting potential cardiotoxicity [56]. Overall, these findings suggest that either the predicted data might inaccurately represent cardiotoxicity as related to Myh7 expression, or there are some potentially poorly understood effects of these chemicals on the heart at high doses, which could arise through direct or indirect mechanisms.

#### 5.4.4 Intestine

When the intestinal lining is damaged transcription up-regulation of various inflammatory signaling genes occurs [70]. One of the genes that is a universal indicator of inflammation is Lcn2 [4]. In DrugMatrix Intestinal gene expression was only measured using the Codelink RU1 arrays and only limited number of treatments were assessed making prediction of gene expression particularly challenging. In the measured data there were a limited number of chemicals that caused up-regulation of Lcn2. These include bisacodyl, loperamide, and azasetron (see Complete DrugMatrix application). At the high doses that were used to create the data in DrugMatrix these chemicals are all plausibly toxic to the intestines.

#### 5.4.5 Kidney

One of the most well-established transcriptional biomarkers of toxicity is Havcr1 (also known as Kim-1). It is induced in the kidney following acute damage [98]. Havcr1 is present on both the Codelink RU1 and the Affymetrix RG230 platforms, with more solely predicted data available for RG230. A review of the measured data reveals an up-regulation of Havcr1 in the kidney following exposure to several well-established nephrotoxic treatments that cause histopathological effects in the proximal tubule.

Examining the purely predicted treatments for kidney that show the most significant up-regulation, we find treatments like cisplatin (28 days, 0.05 mg/kg), bacitracin (3 days, 380 mg/kg), and allopurinol (1-day, 3-day, or 5-day, all at 175 mg/kg) (see Complete DrugMatrix application). Some intriguing predictions also include an up-regulation of Havcr1 by isoprenaline (3 days, 15 mg/kg), thioacetamide (5 days, 200 mg/kg), and sildenafil (5 days, 14.6 mg/kg) (see Complete DrugMatrix application).

Isoprenaline, at high doses, can reduce the glomular filtration rate due to an increase in pre and postglomerular resistance [28], thereby damaging the kidneys [120]. Thioacetamide, while being a reactive chemical primarily known for its liver effects, can also be toxic to the kidneys at high doses [13]. Sildenafil may induce hypotension, leading to kidney hypoperfusion and subsequent hypoxia, similar to the effects seen with non-steroidal anti-inflammatory drugs [68]. Notably, several other NSAIDs are also predicted to elevate Havcr1 expression.

#### 5.4.6 Liver

Gpnmb is a transcriptional biomarker that is up-regulated in response to overt cytotoxicity in several tissues, including the liver [40]. It is known to modulate inflammation through its effects on macrophages [92]. In DrugMatrix, Gpnmb is only measured on the RG230 platform, so the data here solely reflect expression from this platform. The chemicals that elicited the strongest response in the measured data were miconazole, n,n-dimethylformamide, and carbon tetrachloride (see Complete DrugMatrix application). All of these are recognized liver toxicants and demonstrated increased liver enzyme levels in DrugMatrix. Notably, when these treatments were excluded from the training data and subsequently predicted, they showed strong up-regulation of Gpnmb, reinforcing the accuracy of the predictive model.

Chemicals displaying the most significant up-regulation of Gpnmb in the purely predicted data (i.e., without any corresponding training data) include doxapram, carbon tetrachloride, thioacetamide, pralidoxime chloride, and atorvastatin (see Complete DrugMatrix application). All these treatments revealed various forms of liver histopathology in the DrugMatrix database, with some eliciting liver necrosis. This observation suggests that up-regulation of Gpnmb in the purely predicted data is indicative of overt liver damage.

#### 5.4.7 Spleen

The spleen and bone marrow often demonstrate correlated toxicological manifestations. This is because toxicity to the bone marrow can induce stress in the spleen, which then needs to remove damaged blood cells originating from the marrow. In DrugMatrix, both tissues were characterized exclusively using the RU1 platform, thus the predicted data for these tissues is limited to RU1.

The gene Cdkn1a, also known as p21, serves as an indicator of genotoxic damage in both the bone marrow (as discussed earlier in the bone marrow section) and, secondarily, the spleen [111]. The measured treatments that resulted in increased expression of Cdkn1a in the spleen are doxorubicin, mitomycin c, carmustine, leflunomide, and chlorambucil(see Complete DrugMatrix application). All these treatments are either direct-acting genotoxic chemotherapeutics or anti-metabolite therapeutics [102], making the findings of Cdkn1a up-regulation consistent with the genotoxic impacts these agents have on spleen cells. Leflunomide, a pyrimidine synthesis inhibitor, is used as an immune suppressant [35]. It causes lymphoid cellular depletion in the spleen by inhibiting DNA synthesis. Such inhibition often triggers a response from the p53 signaling pathway [46], making the observed up-regulation of Cdkn1a with this treatment expected.

When the data for these treatments were excluded from the model and then predicted, all showed increased expression of Cdkn1a, suggesting the model effectively predicts expression indicative of genotoxic effects in the spleen. In instances where spleen expression data was solely predicted (i.e., no corresponding measured data was available), the chemicals projected to induce the most substantial up-regulation of Cdkn1a were daunorubicin, cyclopropane carboxylic acid, etoposide, doxorubicin (at a different dose level than the measured data), thioacetamide, and doxifluridine (see Complete DrugMatrix application). Daunorubicin, etoposide, doxorubicin, and doxifluridine are all recognized genotoxic chemotherapeutics that induce splenic toxicity secondary to bone marrow toxicity [38, 111]. Cyclopropane carboxylic acid resulted in a decreased leukocyte count and diminished lymphocytes in DrugMatrix, consistent with a chemical causing bone marrow toxicity. Thioacetamide has also been linked with spleen toxicity, likely secondary to its effects on the bone marrow [5].

#### 5.4.8 Skeletal Muscle

Increased expression of the Arrdc2 gene has been linked with chemical-induced cellular injury in skeletal muscle, specifically muscle atrophy [114]. In DrugMatrix, Arrdc2 expression was measured using both the RU1 and R230 platforms. For this analysis, we will focus on the RU1 platform due to its more comprehensive alignment between gene expression and histopathology within DrugMatrix. In the measured data, treatments resulting in the most significant up-regulation of Arrdc2 include simvastatin, pravastatin, fluvastatin, cerivastatin, and lovastatin (see Complete DrugMatrix application). All of these chemicals also demonstrated varying degrees of skeletal muscle toxicity, as evidenced by the histopathology recorded in the DrugMatrix database. Furthermore, they are known to cause rhabdomyolysis [75]. When these treatments were excluded from the training dataset and subsequently predicted, all exhibited the anticipated high level of Arrdc2 expression. This suggests the model is adept at predicting Arrdc2 expression.

Treatments that were purely predicted and exhibited the strongest up-regulation of Arrdc2 include 1-naphthyl isocyanate, omeprazole, tetracycline, chlorpromazine, and lansoprazole (see Complete DrugMatrix application). Out of the chemicals predicted to significantly induce Arrdc2, three can cause rhabdomyolysis. These are the proton pump inhibitors (omeprazole and lansoprazole) [104] and chlorpromazine [106]. The remaining two treatments, 1-naphthyl isocyanate and tetracycline, are not known to cause skeletal muscle damage. As such, their predictions might indicate unreliable Arrdc2 expression in relation to skeletal myotoxicity.

### 5.5 Predicted Apical Endpoint Characterization

A component of the original sparse matrix of measured data encompassed hematology (# endpoints), clinical chemistry (# endpoints), and histopathology data (dose group severity for # pathologies across # tissues). In the current version of the matrix completion, histopathological predictions tend to be overestimated due to targeted histological assessment (i.e., in DrugMatrix histopathology was not performed unless there was an expectation of effect) (see Complete DrugMatrix application). This discrepancy is evident in Table 20, where the measured assessment of histopathology from a 1-naphthylisocyanate-60mg/kg-7day exposure identified 4 effects with an average severity score greater than 1. In contrast, the prediction histopathology indicated 27 effects with an average severity exceeding 1. Notably, the four histopathological outcomes identified in the measured data were correspondingly ranked at the top of the predicted findings list. The potential value of these histopathology predictions lies in using the most extreme cases (i.e., those with the highest predicted average severity) to formulate hypotheses regarding potential histopathological effects. This is because predictions with high severity seem to somewhat align with observed changes. Nevertheless, the overall prediction accuracy for the apical endpoints was suboptimal, indicating the model requires significant refinements to enhance its predictive capabilities.

**Table 20:**
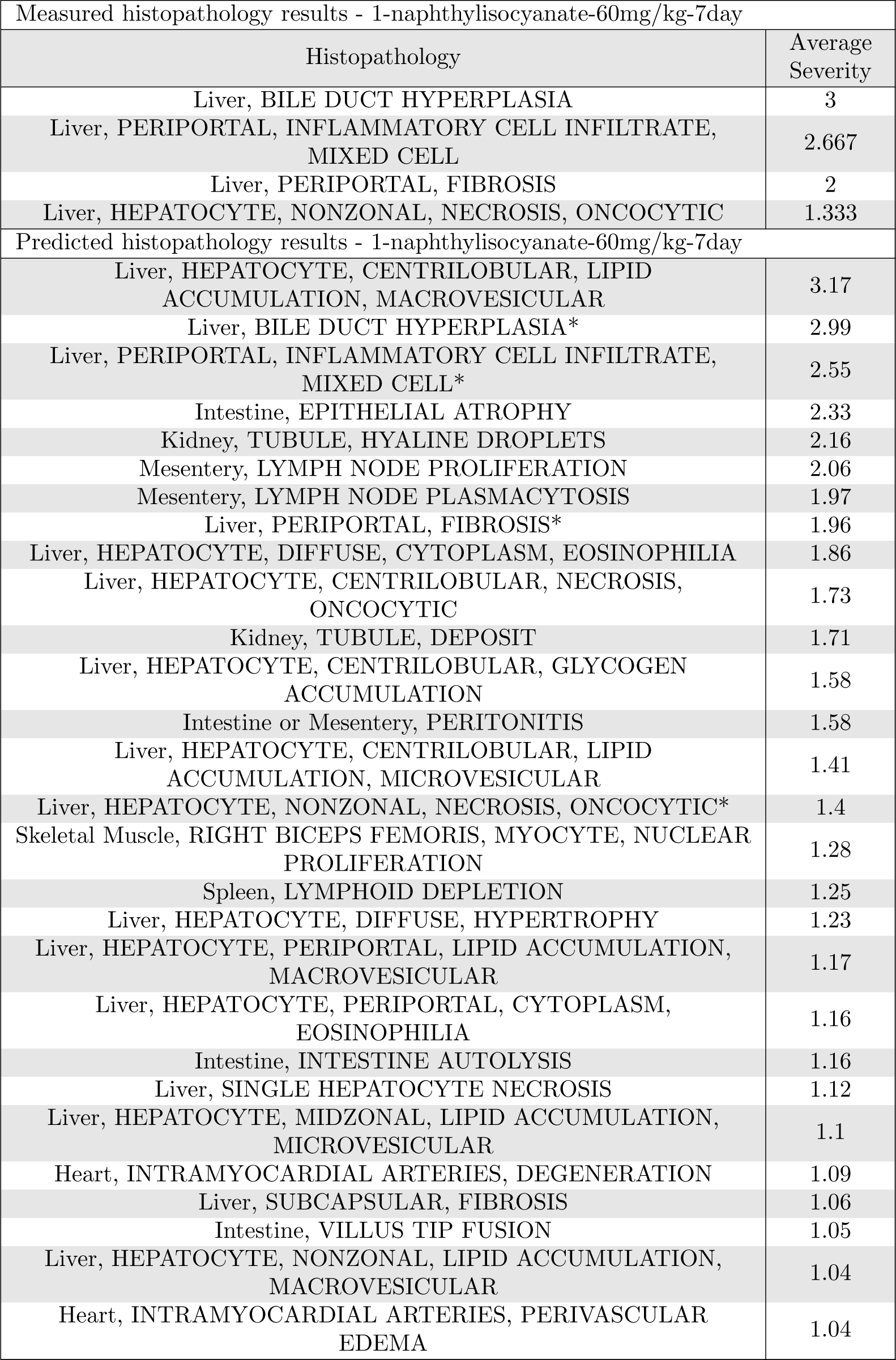
Histopathology, measured vs predicted, 1-naphthylisocyanate-60mg/kg-7day. * Indicates agreement with measure data.

For clinical chemistry and hematology values, the predicted data aligns closely with the measured data (see Complete DrugMatrix application). An example comparing measured and predicted clinical chemistry results for the same treatment (1-naphthylisocyanate-60mg/kg-7day, as mentioned in the histopathology example) is provided in Table 21. As anticipated, there’s a pronounced up-regulation of liver damage biomarkers (alanine aminotransferase, aspartate aminotransferase, and bilirubin) in both the measured and predicted datasets. Additionally, the predicted data indicates a slight increase in another tissue damage biomarker, lactate dehydrogenase. Both datasets also reflect an increase in cholesterol, which is expected for a compound like 1-naphthylisocyanate that inflicts damage to the biliary tract.

**Table 21:**
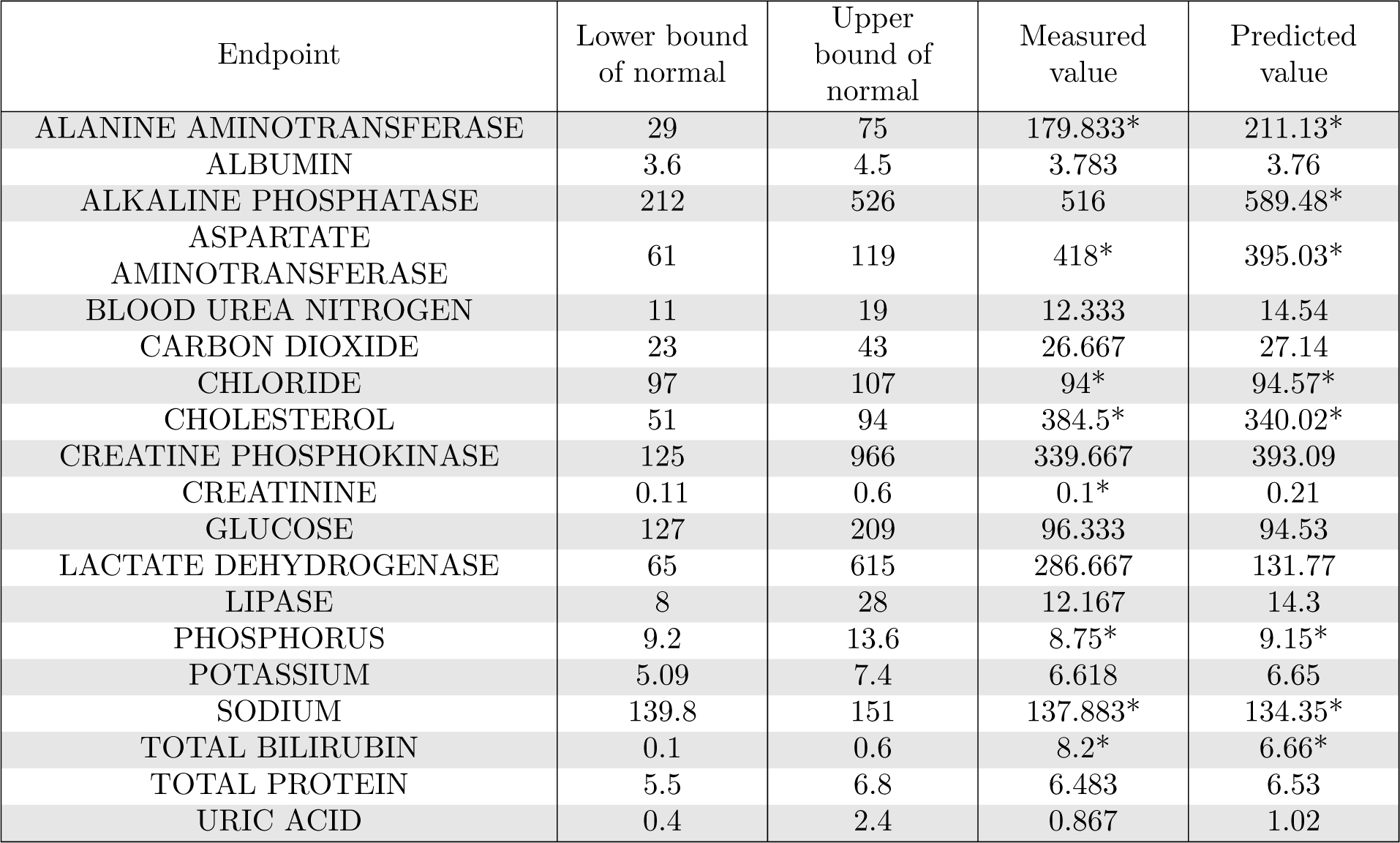
Clinical chemistry, measured vs predicted, 1-naphthylisocyanate-60mg/kg-7day. * Indicates outside of normal bounds.

### 5.6 Web Application for Complete DrugMatrix Access and Analysis

The full predicted DrugMatrix enables further analysis such as clustering drug treatments by their profiles. We performed this clustering and constructed a web application, Complete DrugMatrix, using the R statistical software and Posit’s Shiny framework for R to provide interactive visualizations and enrichment analysis capabilities for these clusters [87, 6, 31, 88, 115, 49, 26, 113, 10, 58, 18, 93, 11, 105, 85, 7, 112]. An instance of the application is available at https://rstudio.niehs.nih.gov/complete_drugmatrix/ and the public code repository is accessible at https://github.com/combspk/Complete-DrugMatrix. We first created separate clusters for each valid combination of tissue and microarray: not all tissues were tested with both the RG230 and RU1 microarrays. For each combination, we constructed a matrix consisting of the log_10_ ratio of treatment gene expression level vs. control gene expression level for each gene for each treatment. A treatment is defined by each combination of chemical, dose, duration, and predicted/experimental status. We then scaled the data in each matrix by calculating the direction-adjusted rank for each gene across each row by first assigning the top 100 genes with the largest absolute value of log_10_ ratios in a row a score from 100 (largest) to 1 (smallest) and every other gene 0. We then multiplied each gene’s ranking by the directionality of its original log_10_ ratio (positive or negative) to get the direction-adjusted rank. If two or more genes had the same log_10_ ratio, they were given the same ranking to have a maximum of 100 ranked genes; that is, if two genes are assigned rank 2, the next highest-ranked gene is assigned rank 4. After scaling the data, we performed dimension reduction using the Uniform Manifold Approximation and Projection (UMAP) package for Python to reduce the matrix to two coordinates per treatment for easy 2-dimensional visualization [73].

The application allows a user to explore the gene expression, histopathology, clinical chemistry, and hematology data for each treatment, if available. Users may search for data for specific treatments by specifying the chemical, dose, duration, tissue, microarray platform, and/or gene name. Users may also opt to see a treatment’s measured data, predicted data, or both. The application also enables users to map the rat genes from each treatment to corresponding human genes using the RG230 and RU1 microarrays and enrich those genes using the Enrichr gene enrichment analysis tool at https://maayanlab.cloud/Enrichr/ [19, 61, 116]. Specifically, Complete DrugMatrix leverages the Enrichr tool to enrich the genes using human-relevant toxicological annotations from the KEGG (2021), BioPlanet (2019), Reactome (2022), and Gene Ontology (2023) databases [45, 53, 51, 52, 39, 25, 9].

Finally, Complete DrugMatrix provides multiple ways to visualize the treatment data and their associated annotations. Figure 9 shows an interactive cluster graph for the *liver, RG230* cluster of treatments. This particular visualization highlights the related chemicals of Fenofibrate, Clofibric Acid, Clofibrate, Gemfibrozil and Pirinixic Acid. We can see that treatments using these chemicals clustered closely together relative to the rest of the *liver, RG230* data based on similar gene expression responses. Furthermore, the application allows a user to view tabular visualizations for the gene expression (see Tables 22, 23), histopathology, clinical chemistry, and hematology data for each treatment. This example shows liver gene expression data for Fenofibrate at both 230 and 430 mg/kg for 5 days. The results show notable correspondence between the measured and predicted data.

**Figure 9:**
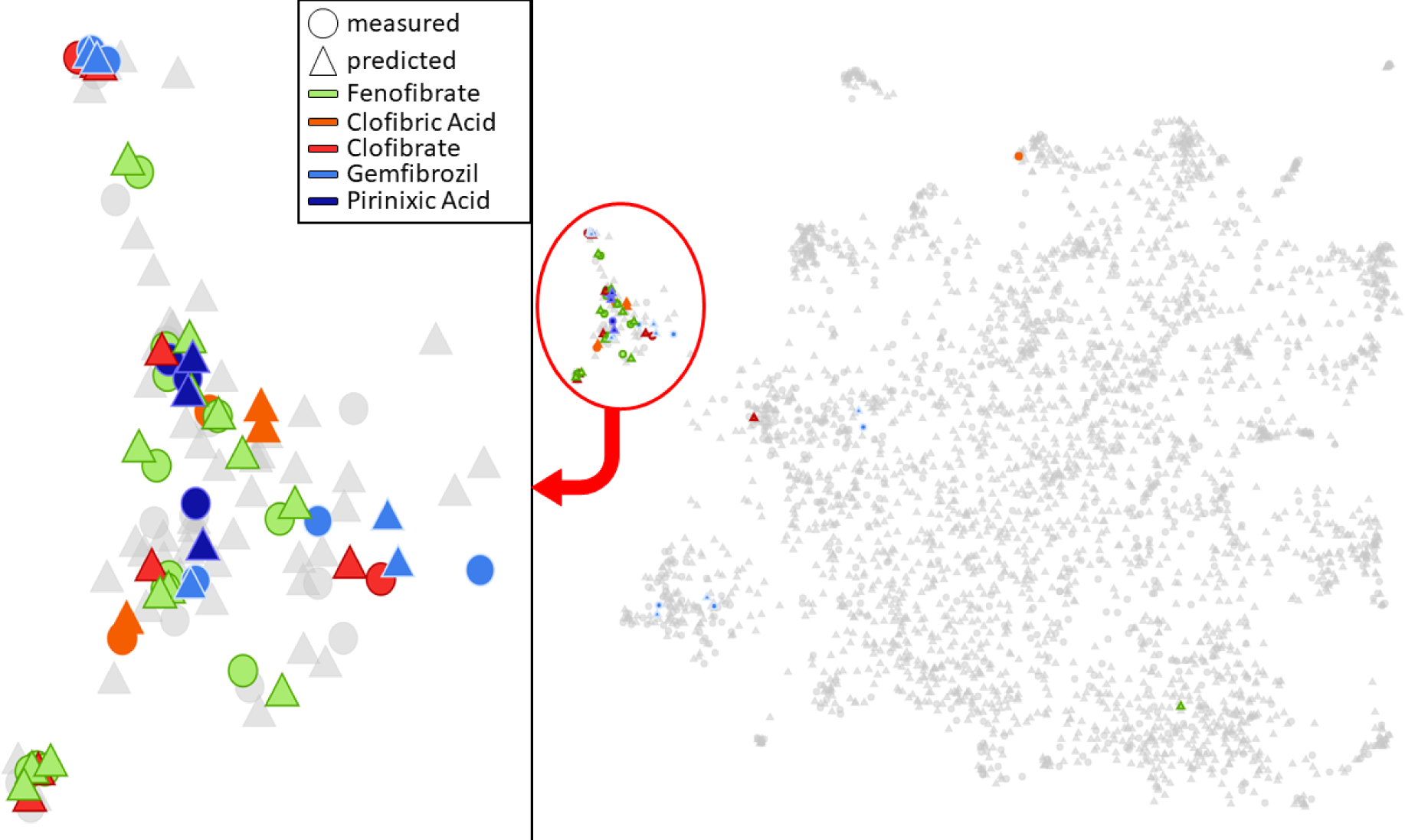
Visualization for the dimension-reduced *liver, RG230* cluster and to be judicious in the application of the data when attempting to draw conclusions related to toxicity, in particular histopathology predictions which as noted in the validation currently over-predict corresponding measured findings.

**Table 22:** Top 10 measured, up-regulated and down-regulated, liver gene expression data (value = log10 ratio treated to control comparison) for Fenofibrate at high doses for 5 days on the RG230 chip pulled from the Shiny application.

| Measured gene expression data for Fenofibrate using RG230, liver mapping |  |  |  |  |  |  |  |  |  |  |
| --- | --- | --- | --- | --- | --- | --- | --- | --- | --- | --- |
| Probeset ID | Human gene | Rat gene | Chip | Tissue | Chemical name | Time | Time unit | Dose | Dose unit | Value |
| P006001 | ACOT1 | Acot1 | RG230 | LIVER | FENOFIBRATE | 5 | d | 215 | mg/kg | 3.218 |
| P006001 | ACOT1 | Acot1 | RG230 | LIVER | FENOFIBRATE | 5 | d | 430 | mg/kg | 3.131 |
| P001780 | AQP3 | Aqp3 | RG230 | LIVER | FENOFIBRATE | 5 | d | 215 | mg/kg | 2.533 |
| P001780 | AQP3 | Aqp3 | RG230 | LIVER | FENOFIBRATE | 5 | d | 430 | mg/kg | 2.244 |
| P018341 | ACOT1 | Acot1 | RG230 | LIVER | FENOFIBRATE | 5 | d | 215 | mg/kg | 2.243 |
| P018341 | ACOT2 | Acot2 | RG230 | LIVER | FENOFIBRATE | 5 | d | 215 | mg/kg | 2.243 |
| P013770 | HDC | Hdc | RG230 | LIVER | FENOFIBRATE | 5 | d | 215 | mg/kg | 2.21 |
| P018341 | ACOT1 | Acot1 | RG230 | LIVER | FENOFIBRATE | 5 | d | 430 | mg/kg | 2.072 |
| P018341 | ACOT2 | Acot2 | RG230 | LIVER | FENOFIBRATE | 5 | d | 430 | mg/kg | 2.072 |
| P013770 | HDC | Hdc | RG230 | LIVER | FENOFIBRATE | 5 | d | 430 | mg/kg | 1.979 |
| ... |  |  |  |  |  |  |  |  |  |  |
| P012822 | RTN4RL2 | Rtn4rl2 | RG230 | LIVER | FENOFIBRATE | 5 | d | 215 | mg/kg | -1.392 |
| P005968 | CYP2C9 | Cyp2c11 | RG230 | LIVER | FENOFIBRATE | 5 | d | 215 | mg/kg | -1.406 |
| P013602 | TSKU | Tsku | RG230 | LIVER | FENOFIBRATE | 5 | d | 215 | mg/kg | -1.421 |
| P027287 | SLC6A6 | Slc6a6 | RG230 | LIVER | FENOFIBRATE | 5 | d | 430 | mg/kg | -1.44 |
| P005891 | CYP2C18 | Cyp2c7 | RG230 | LIVER | FENOFIBRATE | 5 | d | 430 | mg/kg | -1.578 |
| P001730 | APOA4 | Apoa4 | RG230 | LIVER | FENOFIBRATE | 5 | d | 430 | mg/kg | -1.649 |
| P001730 | APOA4 | Apoa4 | RG230 | LIVER | FENOFIBRATE | 5 | d | 215 | mg/kg | -1.665 |
| P005891 | CYP2C18 | Cyp2c7 | RG230 | LIVER | FENOFIBRATE | 5 | d | 215 | mg/kg | -1.714 |
| P006874 | DHRS7 | Dhrs7l1 | RG230 | LIVER | FENOFIBRATE | 5 | d | 215 | mg/kg | -1.906 |
| P006874 | DHRS7 | Dhrs7l1 | RG230 | LIVER | FENOFIBRATE | 5 | d | 430 | mg/kg | -2.657 |

**Table 23:**
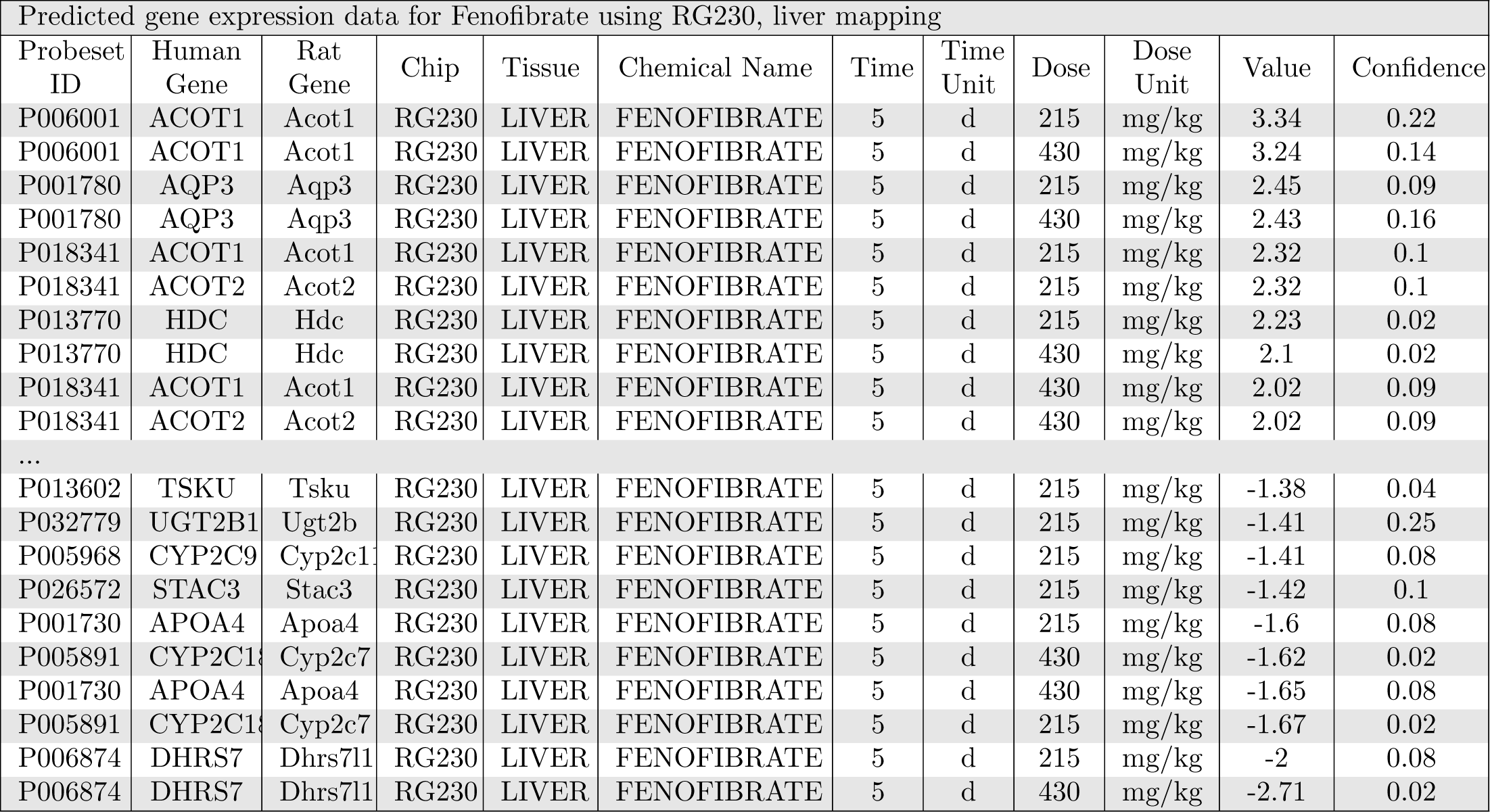
Top 10 predicted, up-regulated and down-regulated, liver gene expression data (value = log10 ratio treated to control comparison) for Fenofibrate at high doses for 5 days on the RG230 chip pulled from the Shiny application.

## 6 Conclusion and Future Work

The DrugMatrix Database represents major effort to create paired traditional and toxicogenomics data to help assess and model toxicity. It is a major resource for the community and used extensively in research and studies. However, DrugMatrix itself is not complete. We present ToxiCompl, an AI method to predict the missing values in DrugMatrix. We show that although vanilla matrix completion yields good MAEs, due to the imbalanced distribution of data in DrugMatrix, important rare signals are lost. To mitigate this issue we formulated a hybrid sampling approach. As a result, ToxiCompl achieves good predictive performance and at the same time retains important rare signals.

DrugMatrix comes with a rich collection of auxiliary data in addition to the matrix itself, and we also explore leveraging such information as side features for two typical machine learning regressors, random forest and deep neural networks. We also implement matrix completion with side features. Our experiments show that random forest and deep neural network regressors do not perform as well as matrix completion through low rank approximation.

We conduct rigorous analysis and validation of predicted data from a biology and toxicology perspective. Through analyzing the connectivity pattern of predicted gene expressions, conducting pathway analysis to characterize molecular pathway level responses from sets of differentially expression genes, leveraging known transcriptional biomarkers of tissue toxicity for validation, and investigating predicted apical endpoint characterization, we show ToxiCompl recovers convincingly many of the missing data that may be useful for formulation of hypothesis around biological effects and toxicological mechanisms a number of unstudied tissues. As ToxiCompl is purely computational and does not require further animal testing, its success points to the potential value of inferential methods that leverage large data sets to predicted complex biological effects of chemicals. Despite the promise of the approach we do caution the end user of the predicted data

Our study and results point to many future research directions. There are several other variants of completion algorithms, for example, factorization machines, completion with cold starts, a hybrid approach of deep neural network with matrix factorization, and graph neural networks that should be further evaluated. In our study, the side features are very simple. With DrugMatrix, we can also leverage the SMILES representation, various finger prints, and the three-dimensional structures of the drugs. Such information obviously makes it possible to design more elaborate features and might improve prediction performance. It is also possible to predict the toxicogenomics profiles for new treatments with zero measurement data.

As was illustrated in the validation, depending on the distribution of the available entries in the matrix (i.e., the measured data), it is not surprising that some predicted entries may be closer to measurement than others. Accurately quantifying the uncertainties and errors in the predicted matrix is essential to practitioners utilizing the predicted DrugMatrix to formulate research hypothesis. Further refinement of the uncertainty metrics will be helpful in the regard and will help the end user understand the limits of applicability of the inferred data.

Complete DrugMatrix has a number of potential uses including 1) inter-tissue response relationships to drug treatment 2) identification of pharmacological signatures across different tissues, 3) building machine learning models to predict the gene responses from a drug treatment and 4) infer a chemical structures that will elicit a specific gene response profiles.

## 7 Acknowledgements

This research used resources of the Compute and Data Environment for Science (CADES) at the Oak Ridge National Laboratory, which is supported by the Office of Science of the U.S. Department of Energy under Contract No. DE-AC05-00OR22725. We would also like to acknowledge Mary Wolfe and Andrew Rooney for implementing and overseeing the interagency agreement between the NIEHS and ORNL that supported the work presented here.

